# Topological basal ganglia model with dopamine-modulated spike-timing-dependent plasticity reproduces reinforcement learning, discriminatory learning, and neuropsychiatric disorders

**DOI:** 10.1101/2025.11.10.687760

**Authors:** Carlos Enrique Gutierrez, Jean Lienard, Benoit Girard, Hidetoshi Urakubo, Kenji Doya

## Abstract

The basal ganglia (BG) are central to action selection and reinforcement learning, yet how the topological organization of the BG circuit with dopamine (DA) D1- and D2-receptors shape learning remains unclear. We present a topologically organized spiking model of macaque BG with cortico-striatal inputs organized into competing channels, D1/D2 medium spiny neurons (MSNs), three-factor DA-modulated STDP for cortical synapses, asymmetric intra-striatal collaterals, and partially overlapping direct/indirect pathways. We validate resting activity and action selection, then study conditioning and generalization–discrimination learning using DA bursts (CS+) and dips (CS).

Two structural determinants emerged. First, pathway overlap (*λ*) trades off selection efficiency and learning speed: higher overlap degrades GPi-based selection efficiency during conditioning, yet accelerates convergence during discrimination by strengthening D2 influence on GPi. Second, lateral inhibition from MSN-D2 to MSNs (*κ*) helps constrain competing actions but is not sufficient alone; robust discrimination requires DA-dip–dependent up-modulation of D2 collateral efficacy (*η*), which speeds and, at low overlap, enables convergence.

Simulations under Parkinsonian and schizophrenia-like settings showed different deficits. A hypodopaminergic “Parkinsonian” STDP regime (D1 LTP loss, D2 LTD loss) impaired conditioning and failed to enhance discrimination. In contrast, attenuated D2 plasticity during DA dips (modeling methamphetamine-induced changes/schizophrenia-related dysregulation) selectively disrupted discrimination while sparing conditioning.

Finally, we demonstrate efficient scaling on the Fugaku supercomputer to rodent and non-human primate–relevant sizes, supporting large-scale, biologically grounded BG simulations. Together, the results highlight how pathway overlap and D2 collateral dynamics jointly regulate the speed and reliability of discrimination learning, and how specific DA perturbations map to distinct learning impairments.

## Introduction

The basal ganglia (BG) are involved in motor and cognitive action-selection and control, with growing evidence supporting their role in reinforcement learning (RL)^1–4^. However, the specific computational processes underlying reward-based learning through spike transmission with different receptors in multiple topological pathways across different nuclei within the BG remain poorly understood.

Biologically constrained models offer a valuable tool for hypothesis testing through simulation^5^ and the exploration of novel theories. Building upon previous works^6,7^, here we develop a topologically organized spiking neural network (SNN) model with synaptic plasticity of the macaque monkey BG. This model incorporates the spatial organization of connections within and across nuclei, distinct subtypes of medium spiny neurons (MSN-D1 and MSN-D2)^8^, dopamine-modulated spike-timing dependent plasticity (STDP) driven by three factors: i) spiking inputs from presynaptic neurons, ii) spiking activity of postsynaptic neurons, and iii) the increase and decrease of dopamine (DA)^9–13^.

The model is based on the anatomical and physiological data from macaque BG^6^, and we investigate the effects of the asymmetry of lateral connections between MSN-D1 and MSN-D2^14,15^ and the overlapping of the MSN-D1 and MSN-D2 outputs to the direct and indirect pathways on RL performance^16–19^.

We first validate the topologically organized BG model by simulating resting state and action-selection under various stimuli.

We then examine the learning performance with the dopamine-modulated STDP in a classical conditioning task with generalization and discrimination learning^13^.

MSN’s axon collateral synapses within the striatum have been suggested to play a potentially critical role in forming and activating functional units^20^, known as channels. Previous experiments have demonstrated that activation of collaterals from MSN-D2 neurons leads to noticeable inhibition of MSN-D1 firing^21^. Our findings suggest that MSN-D2 collaterals are crucial for facilitating discrimination learning.

MSN-D1 and MSN-D2 have different projection targets, with MSN-D1s projecting mainly to the internal globus pallidus (GPi) and substantia nigra pars reticulata (SNr) via the direct pathway, while MSN-D2s project to the external globus pallidus (GPe) via the indirect pathway^22^.

While these pathways are assumed to be segregated in rodents, substantial overlaps have been observed in primates^23,24^. Mounting evidence suggests that direct and indirect pathway neurons cooperate rather than antagonize each other^25^ in controlling movements and that both pathways are necessary for facilitating actions^20,26^. In fact, different degrees of overlaps across pathways have been observed in experiments with mice and monkeys^16–19^ Our work suggests that overlaps of pathways are crucial for discrimination learning and strongly influence the speed of learning.

Finally, we simulate a large-scale version of the BG model on the Fugaku supercomputer, considering realistic sizes ranging from rodents to non-human primates. The simulations employ a hybrid parallelization approach, and computational efficiency is evaluated in terms of network building and simulation time, as well as memory consumption.

## Results

### Topological model of the basal ganglia

A model of the macaque basal ganglia (BG) was previously introduced as a network of point-process neurons lacking spatial structure^7^, with specifications based on anatomical constraints and multi-objective parameter search to match physiological data^6^.

In this study, we incorporate four essential elements to enhance the biological validity of our BG neural circuitry model:

1) The topological organization of neurons, which includes the arrangement of medium spiny neurons (MSNs) as MSN-D1 and MSN-D2 in the striatum, and the organization of synaptic connections within and across the nuclei, with focus on the cortico-striatal afferents as DA-enabled plastic synapses; 2) asymmetric connections between MSN-D1 and D2-MSN-D2 in the striatum; 3) information processing via channels, reflecting the parallel processing capabilities of the BG and shaped by the key anatomical and functional elements underlying channel emergence and organization; and 4) overlapping direct and indirect pathways for experimentation by means of systematic simulation.

These four elements integrated into our model, provide a more comprehensive representation of the BG.

#### 1. Topologically organized Basal Ganglia model

The model incorporates a highly interconnected set of nuclei (Fig.1A): striatal spiny projection neurons or MSNs, fast-spiking interneurons (FSI), subthalamic nucleus (STN), globus pallidus external (GPe), and globus pallidus internal plus substantia nigra pars reticulata (GPi/SNr). Neurons sum up around 27.5 million neurons, where 90 ∼ 96% are MSNs^27,28^. Cortico-striatal (CSN) and pyramidal-tract neurons (PTN) from the cortex^29^ and centromedian/parafascicular neurons (CMPf) from the thalamus are defined as the model inputs, resembling signals from multiple areas converging onto the same area of the striatum^30^. The model parameters, topographical organization and scale are described at methods.

**Figure 1.**
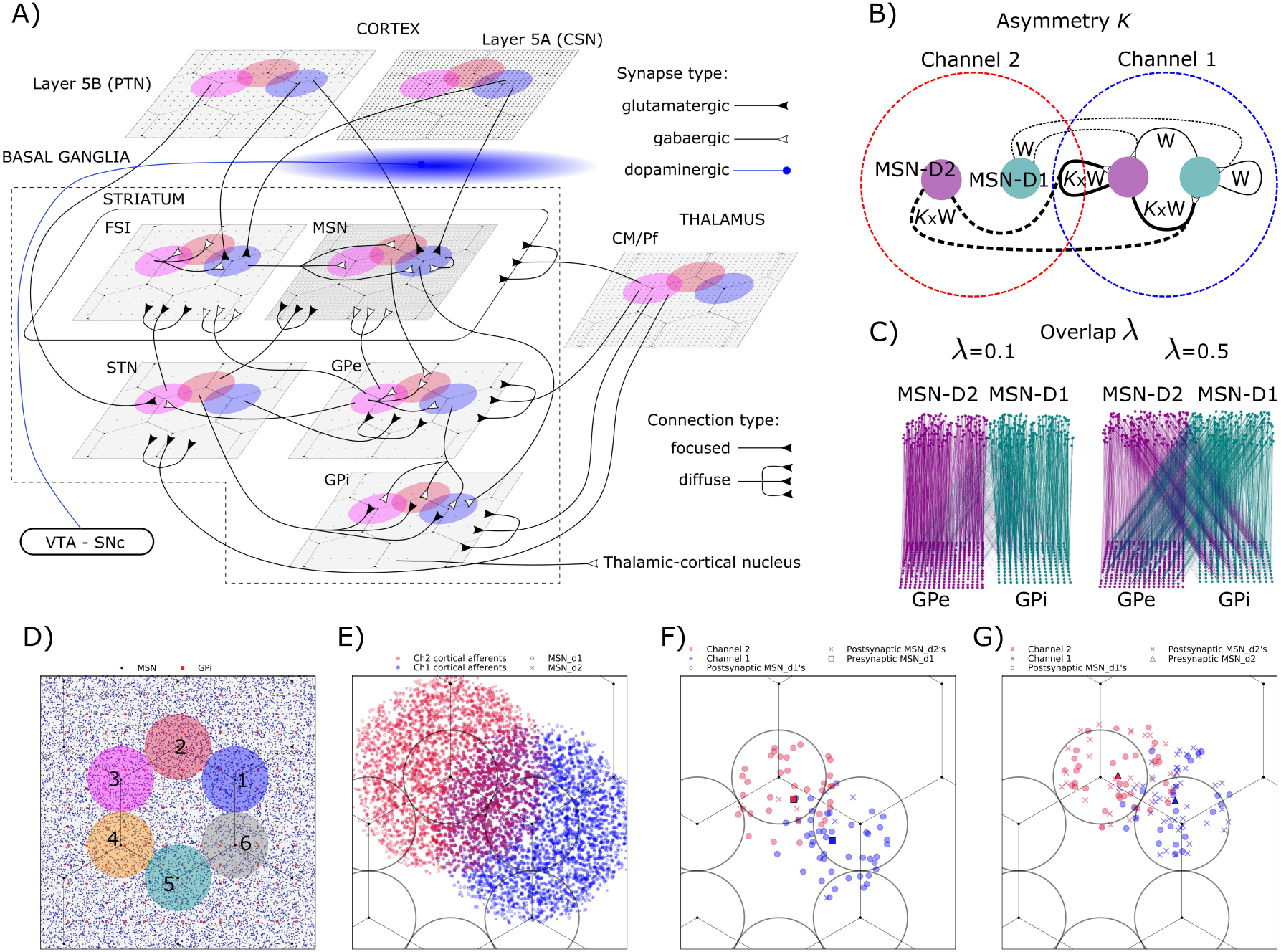
A: schematic figure of the topologically organized model of the basal ganglia. Neurons are allocated spatially across the nuclei by 2D layers with relative positioning. B: schematic figure of two neighboring channels at the striatum and their MSN-D1 and MSN-D2 populations as green and purple circles respectively. At channel 1, the thickness of the connections represents the number of incoming synapses into a single MSN at the target population, where some synapses may come from a neighboring channel 2, indicated with dashed lines. A parameter *κ* is used to explore asymmetric gabaergic collaterals from MSN-D2s. C: the degree of overlap between the direct (MSN-D1->GPi) and indirect (MSN-D2->GPe) pathways is controlled by the parameter *λ* (only few neurons are displayed). A low value of *λ* supports discrimination between pathways, while a high value redirects presynaptic connections between direct to indirect targets. D: the reference 2D surface shows the neural density differences between the input layer (MSN) and output layer (GPi), and six channels of 240*μm* diameter. In dopamine-enabled learning simulations, two adjacent channels were utilized. E: the activation of two adjacent channels by the stimulation of clusters of cortical neurons generates overlapping signals in the striatum and downstream nuclei. F: For two presynaptic MSN-D1 neurons (squares) in different channels, the number of postsynaptic MSN-D1 connections (circles) is similar, while postsynaptic MSN-D2 connections (x’s) are sparse, illustrating a structural asymmetry. G: For two presynaptic MSN-D2 neurons (triangles) in different channels, the number of postsynaptic MSN-D1 (circles) and MSN-D2 (x’s) connections is similar.

#### 2. Striatal asymmetries

Although the influence of connectivity asymmetries between MSNs on dopamine-enabled learning has yet to be fully explored, our model aims to shed light on this area by incorporating two types of asymmetries (Fig.1B):

i. **Structural asymmetry:**^14, 15^ recordings of MSNs indicate that unidirectional connections between MSN-D1 → MSN-D1, MSN-D2 → MSN-D1, and MSN-D2 → MSN-D2 are present in similar proportions, each accounting for approximately 30% of the total observed connections. However, there is a local asymmetry in the number of connections between MSN-D1 → MSN-D2 pairs, which are relatively sparse. In order to model this connectivity in our simulation, the number of incoming connections or indegree parameter *Nsyn* for a single generic MSN is set to 50% of *Nsyn* optimized^7^ for MSN-D1 → MSN-D1, MSN-D2 → MSN-D1, and MSN-D2 → MSN-D2. For MSN-D1 → MSN-D2 connections, we assumed a lower proportion of synapses and set it to 10% of *Nsyn*. This asymmetry is considered a fixed parameter (Fig.1B,F, G). We verified that neglecting up to 40% of *Nsyn* connections from MSN-D1 to MSN-D2 did not impair the model’s ability to reproduce both resting state activity and action selection.
ii. **Strength-dependent asymmetry:** in addition to the proportion of synaptic connections between MSNs, variations in the post-synaptic potentials (PSPs) between them were also observed^14^. Stronger synaptic connections were found between MSN-D2 → MSN-D1,MSN-D2, which may be due to a disparity in the number of GABA receptors. Based on the results presented by Taverna et al. (2008), we estimated that connections with MSN-D2s as presynaptic sources produce PSPs that are at least 2x [mV] in magnitude compared to connections with presynaptic MSN-D1s, and approximately up to 4x [mV]. To model these differences, we assume that the strength of the connections between MSN-D2 → MSN-D1,MSN-D2 is *κ* × *w*, where *w* is the connection strength that is shared by all synapses between MSNs and has been previously optimized, and *κ* (Fig.1B, Fig.4C) is a parameter that represents the degree of asymmetry. Given the potential significance of MSN-D2’s collaterals, we explore the relevance of this asymmetry by varying *κ* between 2.0 and 4.0 and investigating its influence on DA-enabled learning.

#### 3. Channels

The mechanisms behind the formation of parallel information channels in the BG are still not fully understood. However, recent studies using deep brain calcium imaging have identified clusters of MSNs that show synchronized calcium transients during mouse locomotion, suggesting the presence of functional units of processing^31^. These clusters involve both MSN-D1s and MSN-D2s, supporting the idea that MSNs work together to form channels.

Our model incorporates several key features to support the formation of channels.

a. Firstly, the lateral inhibition in the striatum seems necessary for forming channels^20^; thus, gabaergic signals from FSI and MSN’s appear to shape channels.
b. Secondly, the topographical organization of the cortex and its projections to the striatum (Fig.1E, see methods) may support the transmission of discrete cortical inputs within channels across the BG. Channels are believed to be defined by cortical inputs, further emphasizing the importance of cortical-striatal connections in shaping the functional organization of the BG.
c. Thirdly, the topological nuclei connectivity. The model’s focused and diffused type connections (Fig.1A, see methods) may support the synaptic organization of a channel and the spike transmission spread across the nuclei, concentrating synapses on fewer neurons at the output (GPi).
d. Finally, clusters of MSNs appear to be within a circular area of a couple of hundred microns in diameter^31^. Our model assumes adjacent 2D circular regions as relative bounds for channels. This approximation is based on arm-reaching experiments^32,33^, where the spiking activity of eight different channels is associated with the direction to reach. In our model, for the sake of clarity, six channels of 240*μm* in diameter each are allocated in a circular arrangement over the reference patch surface (Fig.1D).

By incorporating these specific features, our model is capable of forming parallel and independent channels that represent different sensory inputs and motor patterns. In simulation experiments, different clusters of cortical neurons were activated to form channels. Each cluster was made up of 200-300 randomly selected neurons from both CSN and PTN within the reference circular area of a channel.

For action-selection and dopamine-enabled learning simulations, two adjacent channels were activated by different cortical inputs, generating overlapping signals in the striatum (Fig.1E) and downstream nuclei. Although the delimitation of channels is unclear, circular reference areas were replicated at relative positions across the nuclei to track the spiking activity of neurons within their scope. The activity of channels can be computed in every nucleus, with the number of neurons per channel described in Table.2. The selected circular area provides a reasonable number of neurons per channel at the GPi and STN for simulations.

#### 4. Overlapping direct and indirect pathways

Recent reports suggest that the traditional view of the BG’s direct and indirect pathways as a dual system may not be entirely accurate. Specifically, axon reconstructions of MSNs have been found to arborize in several areas, including GPi, GPe, and SNr, indicating that the pathways are more complex than previously thought^16,17^. Furthermore, retrograde tracer studies have shown that MSNs project to both GPe and GPi, with collaterals extending to both segments of the pallidum^18^. This overlapping of MSN projections is most prevalent in non-human primates but has also been observed in rodents to a lesser extent^22,34^.

The implications of this convergence and divergence of MSN projections on the BG’s targets cannot be fully understood using traditional approaches to BG organization. To explore the role of these overlapping pathways, our model incorporates a parameter called *λ* (Fig.1C, Fig.4B), which controls the intensity of direct and indirect pathways dichotomy. Simulations with different values of *λ*, ranging from 0 to 0.5, were conducted to analyze the effect of overlap on dopamine-enabled learning.

In the BG model, the indegree parameter was previously optimized to define the number of incoming synapses on every neuron type. A value of *λ* = 0, for example, indicates no overlap, while a value of *λ* = 0.3 means that 70% of incoming MSN’s connections to a single neuron of the GPi/SNr are from MSN-D1s, and 30% from MSN-D2s. Similarly, for a single neuron in the GPe, 70% of the MSN’s synapses originate from MSN-D2s, and 30% come from MSN-D1s.

The striatum receives glutamatergic inputs from the cortex and thalamus, and rewards mediated by dopamine are released diffusely across cortico-striatal projections during learning. Different values of *λ* were tested for a generalization-discrimination learning, and the results suggest that the degree of overlap might be related to the subject’s learning capabilities (see section Overlapping direct-indirect pathways regulate discrimination-learning convergence).

### Baseline state

Before conducting learning tests, the topological BG model was tested to ensure it accurately reproduced the resting state activity and the action selection properties of the previous model^7^.

#### Resting

To evaluate the model’s biological plausibility during the resting state, we compared its network spiking activity during simulation against electrophysiological neural recordings, as in previous studies^7^. The firing rate activities of the model’s inputs were previously optimized to the required levels by adjusting parameters of the corresponding Poisson spike-train sources, one per input population. During resting state simulations, firing rates of 2Hz, 15Hz, and 4Hz were simulated for CSN, PTN, and CMPf, respectively. The resting state simulations were run for 3000ms of biological time, where the first 1000ms simulation time was used for driving the model to steady state and was later excluded from the results. The mean spiking firing rate activity was computed within the [2000, 3000]ms simulation time range. Batches of 10 simulations were run for each combination of the asymmetry *κ* = {2, 4} and the overlap *λ* = {0.2, 0.5}, each using a different random seed per simulation. The nuclei mean firing rates matched the activity of awake monkeys at rest. The simulated baseline activity falls within the biologically plausible values for comparison, previously defined as MSN ∈ [0.01, 1.0], FSI ∈ [7.8, 14.0], STN ∈ [15.2, 22.8], GPe ∈ [55.7, 74.5], and GPi ∈ [59.0, 79.5] (shown as gray areas in Fig.2A). A single simulation spike raster provides qualitative evidence of the results. Exploring parameter values of the asymmetry and overlap showed no effect on the final results, as demonstrated by the mean rate results. Therefore, the topologically organized BG model accurately reproduces the resting state.

**Figure 2.**
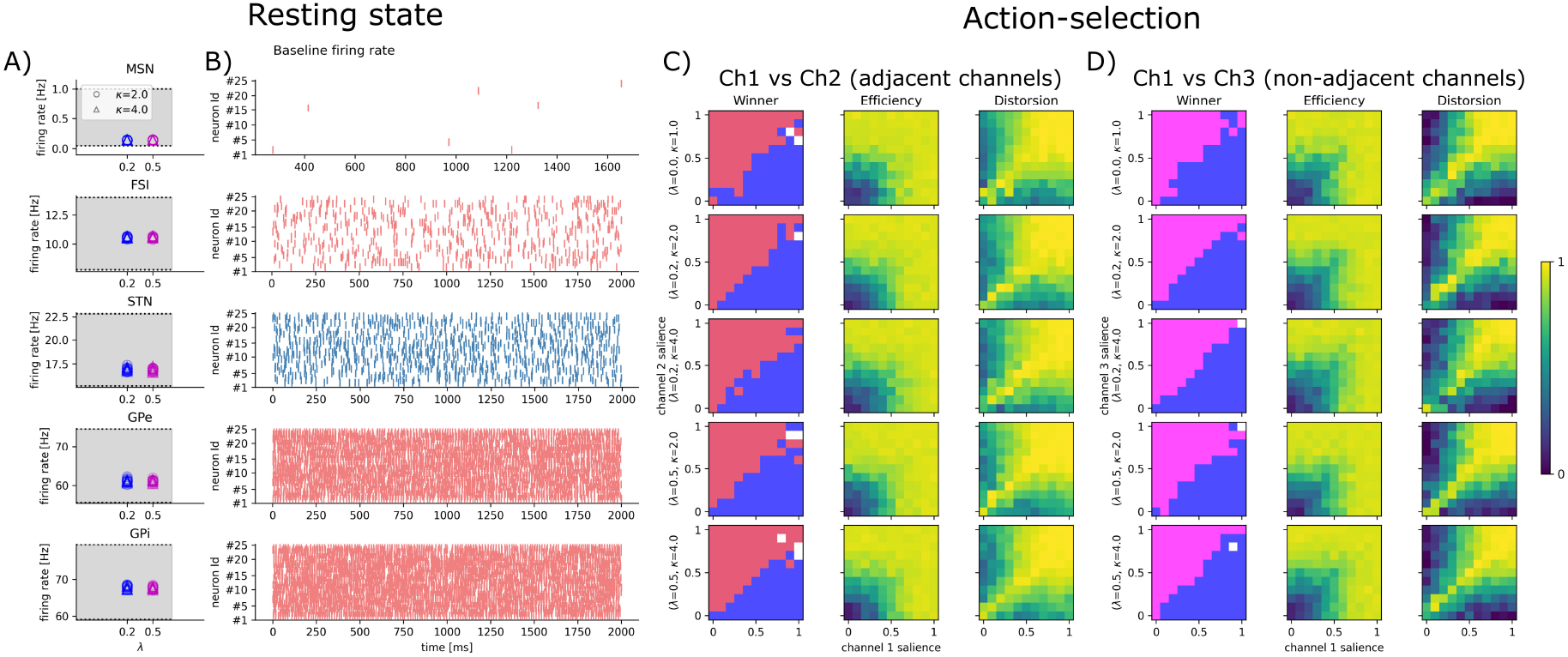
Baseline firing rate and action-selection. A: model baseline activities were evaluated for different combinations of the *κ* and *λ* parameters, with 10 runs for each combination. The resulting mean firing rates fell within the plausible ranges for awake monkeys at rest, as shown by the gray areas. B: qualitatively, the results were confirmed by a single simulation raster plot. C: action-selection assessment by measuring efficiency and distortion matrices for 2 adjacent channels and different values of *κ* and *λ* . D: same as C but for 2 non-adjacent channels. Distortion is lower due to the spatial distance between channels that avoid overlapping signals.

#### Action-selection

To assess the model’s ability to perform action-selection, we systematically stimulated two competitive channels while tracking the lowest GPi/SNr activity, which indicates the selected channel, as in previous studies^7,35^. We evaluated the model’s performance in terms of efficiency and distortion^36^ for pairs of channels, considering values of *κ*={2, 4} and *λ* ={0.2, 0.5}.

We tested adjacent channels with overlapping cortical afferent into their striatal layers, represented by channel 1 (blue) and channel 2 (red), as well as non-adjacent channel 1 (blue) and channel 3 (pink), which resembled information streams corresponding to similar and dissimilar stimuli, respectively (Fig.1D). Each pair of channels received input from three sources: cortical CSN and PTN, and thalamic CMPf. Clusters of 200 PTN cells and 200 CSN cells for each channel, randomly selected from reference circular areas of 240*mμ* diameter, were linearly varied from their baselines of 2 and 15 Hz respectively, to high-activity levels of approximately 20 and 46 Hz, while keeping CMPf at its baseline rate of 4 Hz^7^. The stimuli variations per channel were achieved by activating Poisson spike sources connected to the clusters, whose firing rates gradually increased to fit the required activity levels.

Although neighboring channels 1 and 2 were not clearly dissociated due to overlapping signals from cortical afferents and MSNs axon collaterals, action-selection simulation results showed a relatively high efficiency, indicating the strength of the selection expressed as the lowest GPi channel activity compared with GPi baseline activity. Efficiency increased with the strength of the stimuli, which was similar for both cases of adjacent (Fig.2C) and non-adjacent (Fig.2D) channels .

Similarly, by comparing the lowest GPi channel activity against competitor GPi activity, we measured the distortion in action-selection. Non-adjacent channels displayed less distortion (Fig.2D) than adjacent channels (Fig.2C) due to lower interference from overlapping inputs and collaterals.

However, we observed strong distortion for both cases with competitive stimuli of similar strength, which was noticeable at the anti-diagonal region of the distortion matrices.

Different values of *κ* and *λ* did not produce any relevant difference in efficiency and distortion results. However, this may not be the case when action-selection is performed systematically on dopamine-enabled learning trials (see next sections).

### Generalization-Discrimination Learning

Our aim is to investigate dopamine-enabled learning by simulation of MSN-D1 and MSN-D2 neurons receiving sensory-motor inputs^30^ through CSN and PTN neurons, along with a baseline thalamic input, in a generalization-discrimination learning paradigm.

Recent research has shown that dopamine plays a critical role in regulating generalization-discrimination learning^13^. Specifically, D1-dependent conditioning facilitates learning generalization across similar and unseen sensory stimuli, while subsequent discrimination tasks refine predictions based on dopamine dips detected by D2 receptors when predictions are incorrect. We refer to a correct prediction as the selection of the preferred channel for a given task.

Transient increases in dopamine concentration (DA bursts) enhance the plasticity of dendritic spines in MSN-D1 cells^12^, while transient reductions in dopamine concentration (DA dips) induce spine plasticity in MSN-D2 cells^13^, with both spine enlargements occurring only within a narrow time window after stimulus onset.

In this section, we describe the implementation and simulation of this learning in our topological model, specifically in the context of two neighboring BG channels in competition. We explain the implementation of the learning process as a set of three parts: **conditioning, generalization test, and discrimination**. Next, using systematic simulations of learning episodes, we explore the effects of increasing and decreasing dopamine on different receptors, the changes in spiking activity of MSN-D1 and MSN-D2 cells, and how these changes affect learning. Our simulation results reveal how different degrees of lateral MSN-D2 inhibition *κ* and overlapped direct-indirect pathways *λ* affect the timing of discrimination learning, providing new clues into their functional roles.

#### Conditioning (CS+)

To establish classical conditioning (CS+), sensory **stimuli A** are modeled (inputs from PTN and CSN) and presented on adjacent channels 1 and 2 (Fig.1D), with channel 1 as the preferred channel. Two clusters of 300 PTN and 300 CSN neurons for each channel are randomly selected and activated with spikes generated from a Poison process on each channel, resulting in cortical firing activity. Stimuli A deliver signals with slightly higher amplitudes (firing rates) to the preferred channel than to channel 2 (Fig.3A). Input strengths were matched to those used in the action-selection task. As in that section, CSN and PTN firing rates were varied linearly from baseline (2 and 15 Hz) to high-activity levels (∼20 and ∼46 Hz) across 11 steps (0–10). For Stimulus A, channel 1 corresponds to step 8 (CSN ≈ 16.4 Hz; PTN ≈ 39.8 Hz), and channel 2 to step 6 (CSN ≈ 12.8 Hz; PTN ≈ 33.6 Hz).

**Figure 3.**
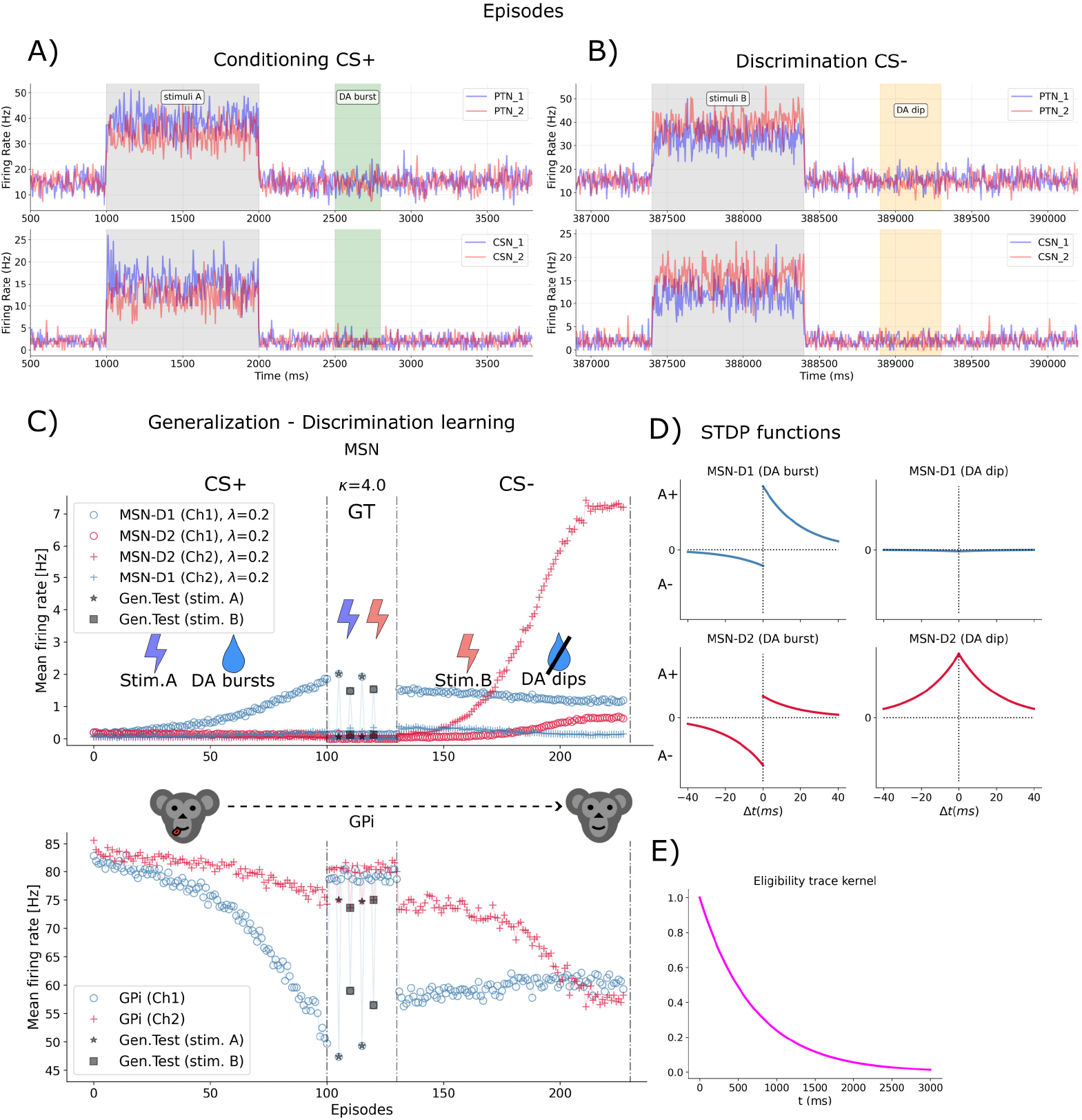
Episodes. A, B: sensory stimuli A and B were modeled based on inputs from PTN and CSN with similar firing rates on adjacent channels 1 and 2. During the conditioning phase (A), stimulus A was slightly higher on channel 1, while during the discrimination phase (B), stimulus B was higher at channel 2. The timing parameters for stimulation, DA burst, and DA dip were derived from experimental results^12,13^. C: Learning stages. CS+: the mean activities of MSN and GPi neurons were computed during stimulation at channels 1 and 2. In the first trials, MSNs are mostly silent while the GPi selected either channel 1 or 2, allowing the model to explore actions. Eventually, the preferred channel 1 was more frequently selected, resulting in DA release as a reward. Across subsequent trials, synaptic plasticity caused an increase in the activity of MSN-D1s at channel 1. GT: to confirm CS+ learning, 2 trials for stimuli A and B each lead to the selection of channel 1, generalizing the model response for similar inputs. CS-: for discrimination learning, we presented sensory stimuli B to both channels. When the learned action (channel 1) was selected, the reward was omitted, triggering a DA dip that promoted plasticity on MSN-D2s. Over episodes, the GPi activity at channel 2 gradually shifted towards channel 1, allowing the model to discriminate between stimuli. D: STDP functions. E: eligibility trace kernel for a time constant parameter *τ*_*c*_ = 700*ms* for allowing plasticity within the critical time window defined in A and B.

In each trial (episode), stimuli A last for 1s of biological time, and the winner channel is determined by computing the lowest mean firing rate from GPi channels 1 and 2 within the 1s stimulation time window, considering the activity of the neurons within the reference circular masks of channels 1 and 2. Upon its selection, the preferred channel is then paired with a US (unconditional stimulus), resulting in a reinforcement signal that is assumed to be a positive reward prediction error^37^, expressed as transient DA increases.

The transient increase in DA (burst) is modeled as a Poisson process that emits spikes at a frequency of 20Hz^38^ from a baseline of 0Hz. We assumed 0Hz to represent the tonic DA activity, in order to avoid changes in the synaptic strengths of cortico-striatal connections due to non-zero baseline cortical activity. This assumption was made in conjunction with the use of the dopamine-based synapse model from the NEST neural simulator^39^ (see methods).

DA burst is simulated after 1.5s from the onset of stimuli A, and lasts for 0.3s, within the critical time window^12^ (Fig.3A). The reinforcement DA signals are diffusely released on all cortico-striatal synapses, regardless of the predefined competing channels.

Following reinforcement, the simulation runs for a period of 1s under baseline conditions until the onset of the next episode. Throughout the early trials, MSNs are almost silent, and the model selects channel 1 or 2 as in a RL exploratory stage. The first batch of trials, approximately the first 10-20 episodes, can be assumed to be a sort of exploration phase.

As training progresses, the synaptic plasticity of MSN-D1 leads to an increase in the excitability of these cells, and the afferent synapses from cortex to MSN-D1, particularly those targeting channel 1, strengthen as a result (Fig.3C). We run a total of 100 episodes for the CS+. For details on the SDTP functions (Fig.3D) please refer to the Methods section.

#### Generalization test (GT)

To verify the success of the CS+ training and test for generalization, we conducted several trials without DA activation. We presented stimuli A or a similar one, stimuli B, to test whether the model could select the preferred channel. Stimuli B is defined symmetrically to stimuli A but with the amplitudes reversed across channels (Fig.3B): it delivers slightly stronger input to channel 2 and slightly weaker input to channel 1. Using the same scaling, channel 2 corresponds to step 8 (CSN ≈ 16.4 Hz; PTN ≈ 39.8 Hz) and channel 1 to step 6 (CSN ≈ 12.8 Hz; PTN ≈ 33.6 Hz).

When presented with stimuli A (which was slightly stronger on channel 1), the model consistently selected the CS+ preferred channel 1, confirming the success of the classical conditioning training. When presented with stimuli B (which was slightly stronger on channel 2 (Fig.3B)), designed for discrimination learning CS-(see next section), the model also selected the previously learned channel 1, indicating that the D1-based conditioning learning had generalized to similar stimuli. We performed two trials each for stimuli A and B to ensure reliable results (Fig.3C). In addition, in the absence of stimuli, MSN and GPi spiking activities return to their baseline values.

#### Discrimination (CS-)

To distinguish between reward-predicting and non-reward-predicting stimuli during generalization, we test whether our model could discriminate between the two. We conducted approximately 100 trials for discrimination learning. During the trials, we presented sensory **stimuli B** to channels 1 and 2. These stimuli are designed with a slightly higher amplitude (firing rate) towards the new preferred channel 2 than channel 1. The stimuli last for 1s of biological time (Fig.3B).

Channels 1 and 2 are stimulated, and when the learned action (channel 1) is selected, the reward is not delivered (CS-). The omission of the expected reward triggers DA dips for 0.4s of biological time after 1.5 seconds from the onset of stimuli B, within the critical time window^13^.

We implemented the plasticity effect of DA dips on cortical afferents targeting MSN-D2 cells using an STDP update rule enabled by spikes from a Poisson process resembling DA neurons. This STDP rule is different from the conditioning learning case (Fig.3D, see methods section).

The NEST neural simulator currently does not support STDP that is driven by transient changes from tonic to lower activities of cells (DA dips). Therefore, we used the same approach as in the CS+ case but targeted different synapses. Specifically, we distinguished synapses on D2-receptors from those on D1-receptors in our model. This allowed us to enable STDP on MSN-D1s or MSN-D2s depending on whether the stimulus was a CS+ or CS-, respectively.

DA dips were also delivered diffusely across all cortico-striatal afferents, which increased the excitability of MSN-D2 cells, especially those at channel 2 over time (Fig.3C). With DA dips, channel 2 activity at the GPi gradually approached channel 1 GPi activity until convergence. This means that the new preferred channel 2 is selected, and DA dips were no longer delivered. The critical period of DA dips, coupled with STDP, led to discrimination between reward-predicting and non-reward-predicting stimuli.

#### Simulations

We systematically explored the impact of different values of the overlap degree parameter *λ* (Fig.4B), which defines the cooperation degree between MSNs on downstream spiking activities (GPe and GPi), thereby affecting the activation of a channel.

**Figure 4.**
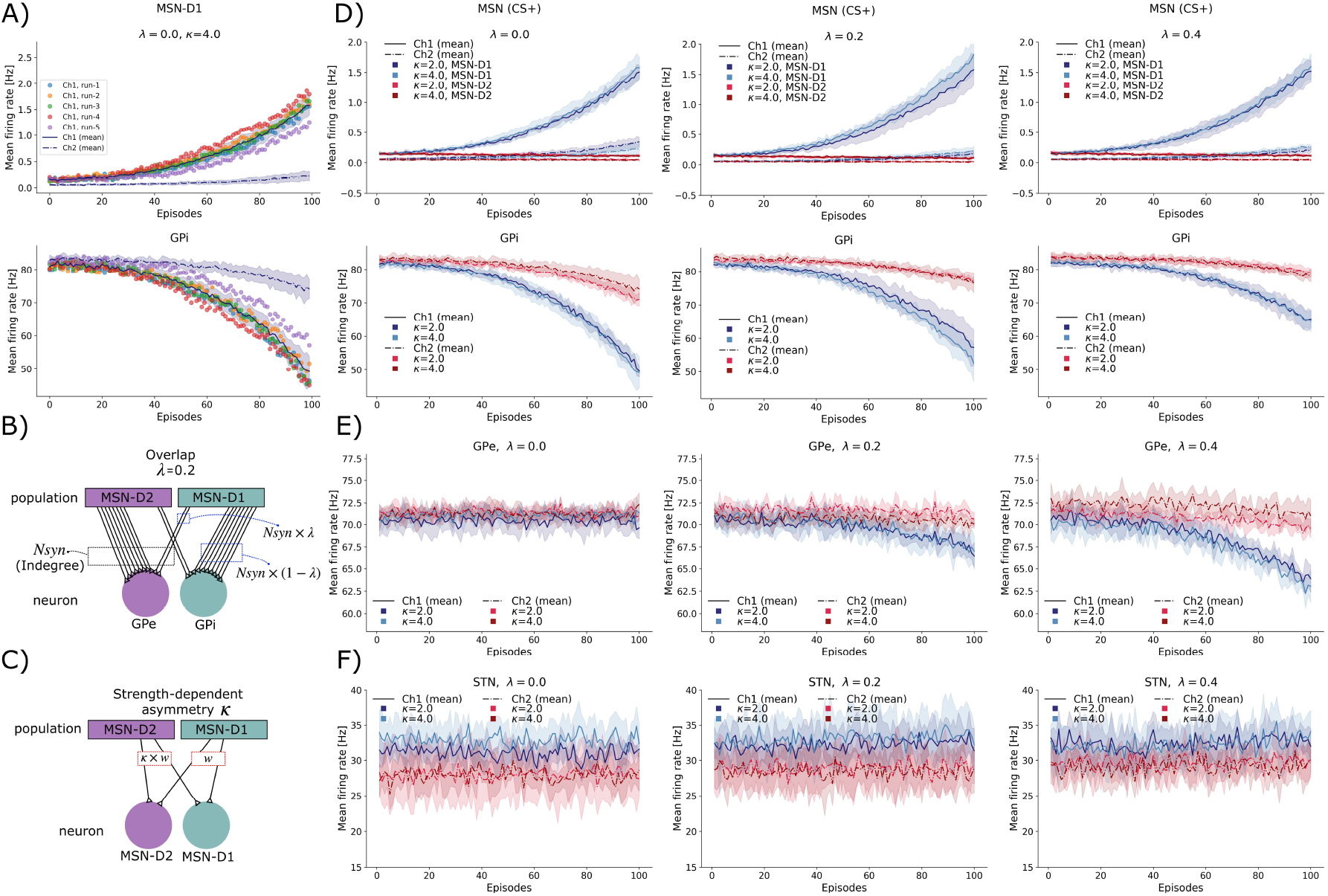
CS+. A: For each parameter combination (*κ* and *λ*), simulations were run for CS+ (n=5), each consisting of 100 episodes with different random seeds. Each dot on the graph represents the mean activity of MSN-D1 neurons (upper panel) and GPi neurons (lower panel) within a channel during a single episode. The results from the five runs were averaged to generate a mean curve. B: Schematic representation of the overlap parameter *λ* = 0.2, where 80% of the incoming synapses (*N*_syn_) to GPi (or GPe) originate from MSN-D1 (or MSN-D2), and the remaining 20% originate from MSN-D2 (or MSN-D1). C: Illustration of the asymmetry parameter which modifies the connection strength (*w*) of synapses from MSN-D2 to other MSNs by a factor of *κ*. D: At channel 1, MSN-D1 neurons exhibit consistent mean firing activity across different *λ* and *κ* values, whereas GPi disinhibition varies with *λ* . E: GPe inhibition at channel 1 increases as *λ* increases. F: In the simulations, GPe inhibition has little effect on STN activity, resulting in no significant changes.

We also varied the strength-dependent asymmetry *κ* (Fig.4C) which changes the collateral transmission of GABA from MSN-D2s to other MSNs during trials of CS+ and CS-. Giving neighboring channels, we may assume that the effect of *κ* on a certain channel would come not only from local MSN-D2s but also from MSN-D2s at the neighboring channel. Collaterals from FSI and MSN-D1’s cells were assumed to be fixed parameters.

For each combination of parameters *λ* and *κ*, we conducted *n* = 5 runs (Fig.4A) of the generalization-discrimination learning^13^ using different random seeds for each run on the NEST neural simulator. The number of episodes for a single run is 230 = (100CS+) + (30GT) + (100CS-).

The spiking activities of cells within channels are summarized as mean firing rates, calculated within the window of time for sensory stimulation. For instance, in Fig.4A, single dots represent mean firing rates of MSN-D1 (top) and GPi (bottom) at channel 1 for different episodes. Runs are summarized as mean curves with standard deviation. These curves represent how the neural activity changes, in average, over episodes for a certain combination of *λ* and *κ*.

### Overlapping direct-indirect pathways decreases action-selection efficiency in conditioning learning

During CS+ episodes, the rate of change in the mean firing rates of MSN-D1s at channel 1 shows consistent patterns across different values of *λ* and *κ* (Fig.4D). This suggests that the explored parameters minimally impact MSN-D1 activity, which is primarily driven by the potentiation of afferent connections from the cortex to MSN-D1s.

For different *κ* values, the inhibitory effect of the collaterals from MSN-D2s to MSN-D1s remains minimal, as MSN-D2 activity stays close to baseline levels at approximately 0 Hz (Fig.4C). However, during CS+ episodes, we observed that GPi disinhibition in channel 1 increases with increasing *λ*, thereby reducing the selection efficiency, which is measured as the distance between GPi channel 1 and GPi baseline activities (Fig.4D).

Higher levels of overlap *λ* alter the number of incoming GABAergic inputs to the GPi from both MSN-D1 and MSN-D2 neurons (Fig.4B). As *λ* increases, the inputs from MSN-D1 neurons decrease, while those from MSN-D2 neurons increase. Since MSN-D2 neurons remain at baseline activity, this shift may help explain the desinhibitory effect observed in GPi activity.

Additionally, increasing *λ* raises the incoming GABAergic inputs from MSN-D1s to the GPe, resulting in increased inhibition of the GPe at channel 1 (Fig.4E). This inhibition intensifies during conditioning learning.

### Influence of collaterals from MSN-D2 during discrimination-learning

After CS+ completion using stimuli A, we tested the generalization of conditioning learning using a similar stimuli B and confirmed the persistent learned selection of channel 1 (Fig.3C). To further refine this prediction towards the selection of channel 2, as suggested by Iino et al.(2020)^13^, stimuli B was presented throughout CS-trials (Fig.5) with varying values of *λ* and *κ*.

**Figure 5.**
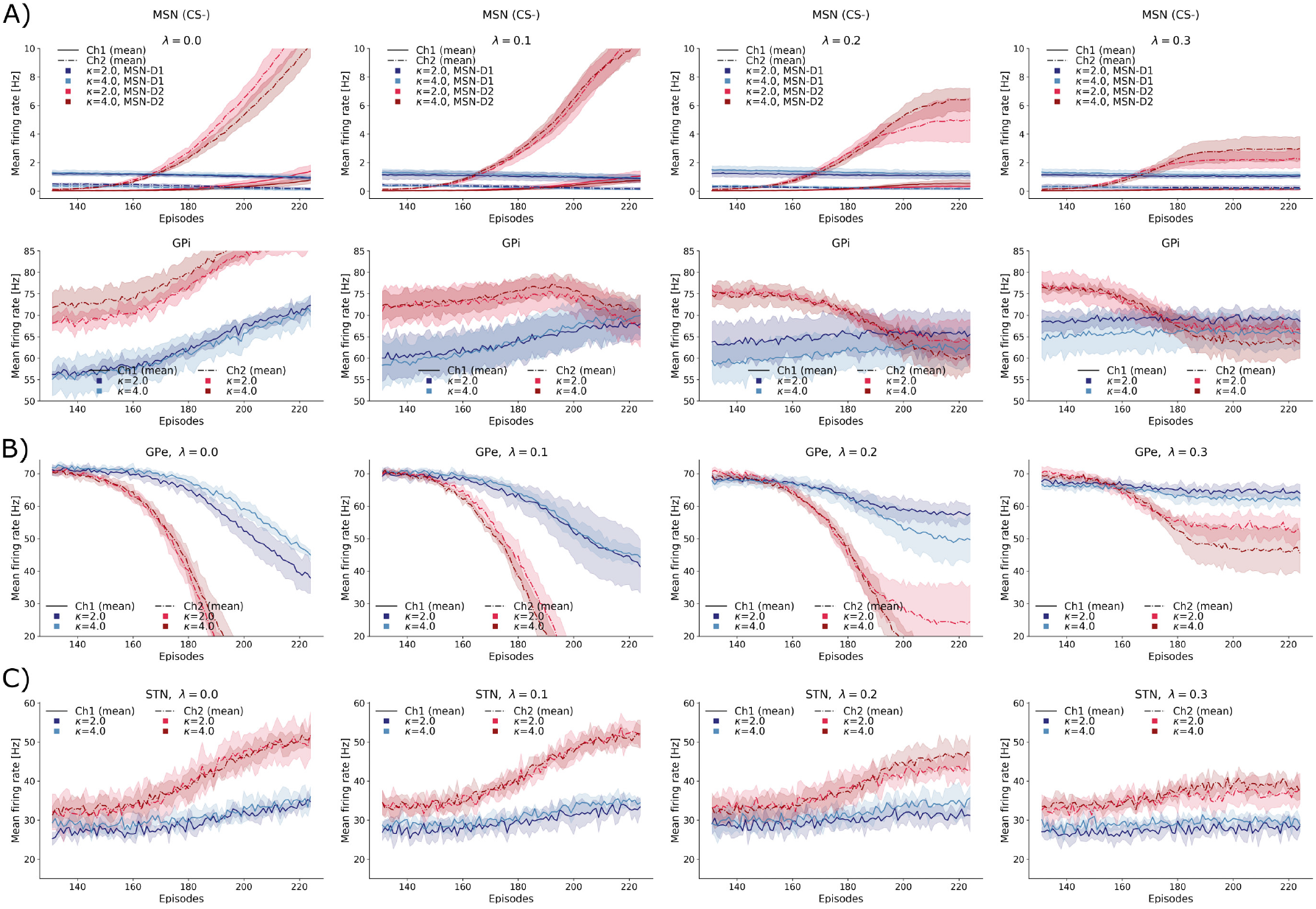
CS-. A: Plasticity enables MSN-D2 neurons at channel 2 to increase their activity independently of *λ* or *κ*, resulting in enhanced lateral inhibition from MSN-D2 to MSN-D1 neurons at channel 1. Over episodes, GPi at channel 1 becomes disinhibited, promoting convergence, while GPi at channel 2 undergoes gradual inhibition, a process that accelerates with higher *λ* values. B: Increased activity of MSN-D2 neurons at channel 2 leads to GPe inhibition and the disinhibition of GPe at channel 1, with faster dynamics observed at higher *λ* values. C: STN activity at channel 2 progressively increases over episodes.

Over the trials, the excitability of MSN-D2 cells at channel 2 increased across all configurations of *λ* and *κ*, but only up to certain trials of CS-. This increase was driven by the plasticity of cortical afferent synapses onto MSN-D2s, enabled by DA dips (Fig. 5A). However, the convergence of MSN-D2 activity was dependent on *λ*, with lower *λ* values leading to higher final firing rates and higher *λ* values resulting in lower final firing rates.

Lateral inhibition with *κ* = 2.0, 4.0 from MSN-D2s at channel 2 influenced the firing activity of MSN-D1s at channel 1, resulting in a gradual decrease in MSN-D1 activity as MSN-D2 firing rates increased over the trials, as shown in the upper panel of Fig.5A. A magnified view highlights the slow changes in MSN-D1 activity at channels 1 and 2 across trials for different *κ* values, under both low (Fig.6A) and high (Fig. 6B) *λ* conditions. Additional collaterals may originate from MSN-D2s at channel 1, which slightly increased activity towards the end of CS-due to the plasticity of afferent synapses from cortex to channel 2, with some overlap on channel 1, and from FSIs.

**Figure 6.**
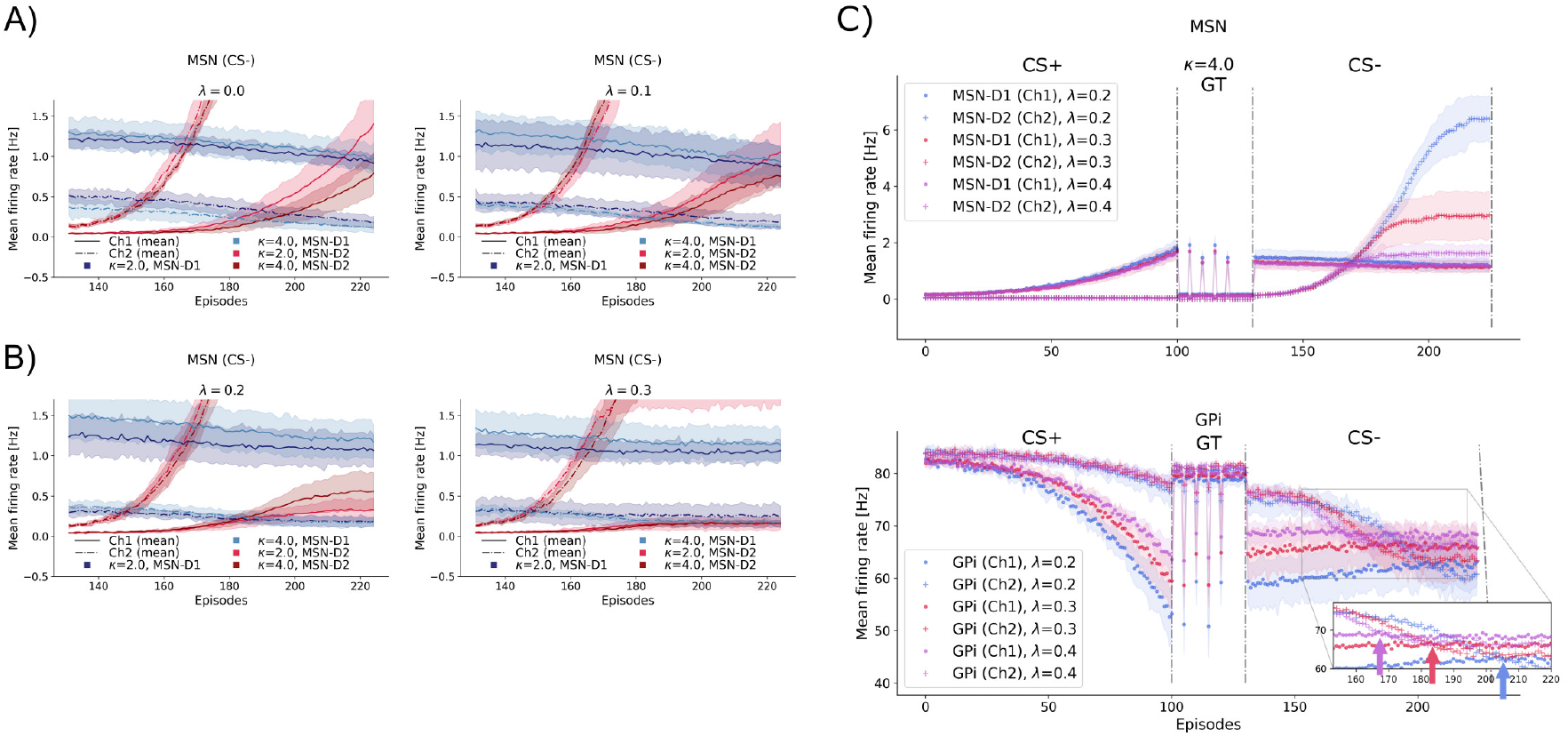
Collaterals during CS-. A, B: Magnified views of the upper panel of Fig.5A, illustrating that lateral inhibition (mediated by *κ*) from MSN-D2 neurons at channel 2 leads to a slight reduction in MSN-D1 neuron firing during CS-episodes under low (A) and high (B) pathway overlap. C: Discrimination learning is modulated by *λ* . Diverse convergence patterns, indicated by arrows, show the selection of channel 2 at the GPi for different *λ* values, with faster learning observed at higher *λ* . This is also reflected in the increased mean firing rate of MSN-D2 neurons at channel 2 during CS-, transitioning to a steady activity level (indicating learning completion) over different episodes.

Collateral projections impact downstream nuclei, which is particularly noticeable in the case of non-overlapping pathways (*λ* = 0). In this scenario, the GPi at channel 1 (Fig. 5A, lower panel) is gradually disinhibited. When pathways are perfectly discriminated, GPi disinhibition at channel 1 may be driven entirely by MSN-D1s, whose activity may be modulated by MSN-D2 collaterals from channel 2. Furthermore, unlike CS+ condition, we observe in CS– condition that GPi disinhibition for channel 1 is slightly stronger at lower *λ* values.

Additional sources of GPi deshinbition may arise from GPe afferents (Fig.5B). GPe firing activity is reduced, along with its inhibitory effect on GPi, due to increased MSN-D2s activity (for *λ* =0, GPe’s inhibition may come entirely from MSN-D2s). Another additional source may be the STN (Fig.5C), which projects diffusely to GPi and increases its excitatory input over CS-.

GPi disinhibition at channel 1 supports the convergence of discrimination learning. Specifically, upon presentation of stimulus B, there is a gradual inhibition of the GPi at channel 2 due to increased activity of MSN-D2s in channel 2. This inhibition continues until the activity of GPi at channel 2 falls below that of GPi at channel 1 (Fig. 5A for *λ >* 0.0), ultimately leading to the selection of channel 2.

In our simulations, significant differences between *κ* = 2.0 and *κ* = 4.0 were not observed.

### Modulation of collaterals from MSN-D2 to MSN, enabled by DA dips, is crucial for achieving convergence in discrimination learning

Our results indicate that MSN-D2 collaterals play a crucial role in the disinhibition of the GPi at channel 1 required for the convergence of discrimination learning during CS-, primarily by modulating the firing rates of MSN-D1s. Additionally, an optimal value of *λ* appears to be required to achieve convergence. For a low degree of pathway overlap, we observed that CS-convergence may not occur or may require a greater number of episodes, which seems biologically implausible and unrealistic.

Recent findings have further illuminated the modulatory effects of DA on axon collaterals within the striatal circuitry. Using transgenic mouse brain slices and optogenetic stimulation, researchers isolated different synaptic connections between MSNs.

Whole-cell recordings revealed that DA activation leads to the depression of GABAergic transmission from presynaptic MSN-D2 neurons, mediated by presynaptic D2 and serotonin 5-HT1B receptors, with D2 receptors accounting for approximately half of the observed depression^40^. Thus, DA bursts enabled a modulatory effect that depresses collaterals from MSN-D2s to other MSNs (Fig.7A). In our simulation, this modulatory effect was not included; however, we found that MSN-D2 collaterals had almost no effect during CS+ (*κ*=2.0, 4.0 at Fig.4D), corresponding to the notion of weak collaterals.

**Figure 7.**
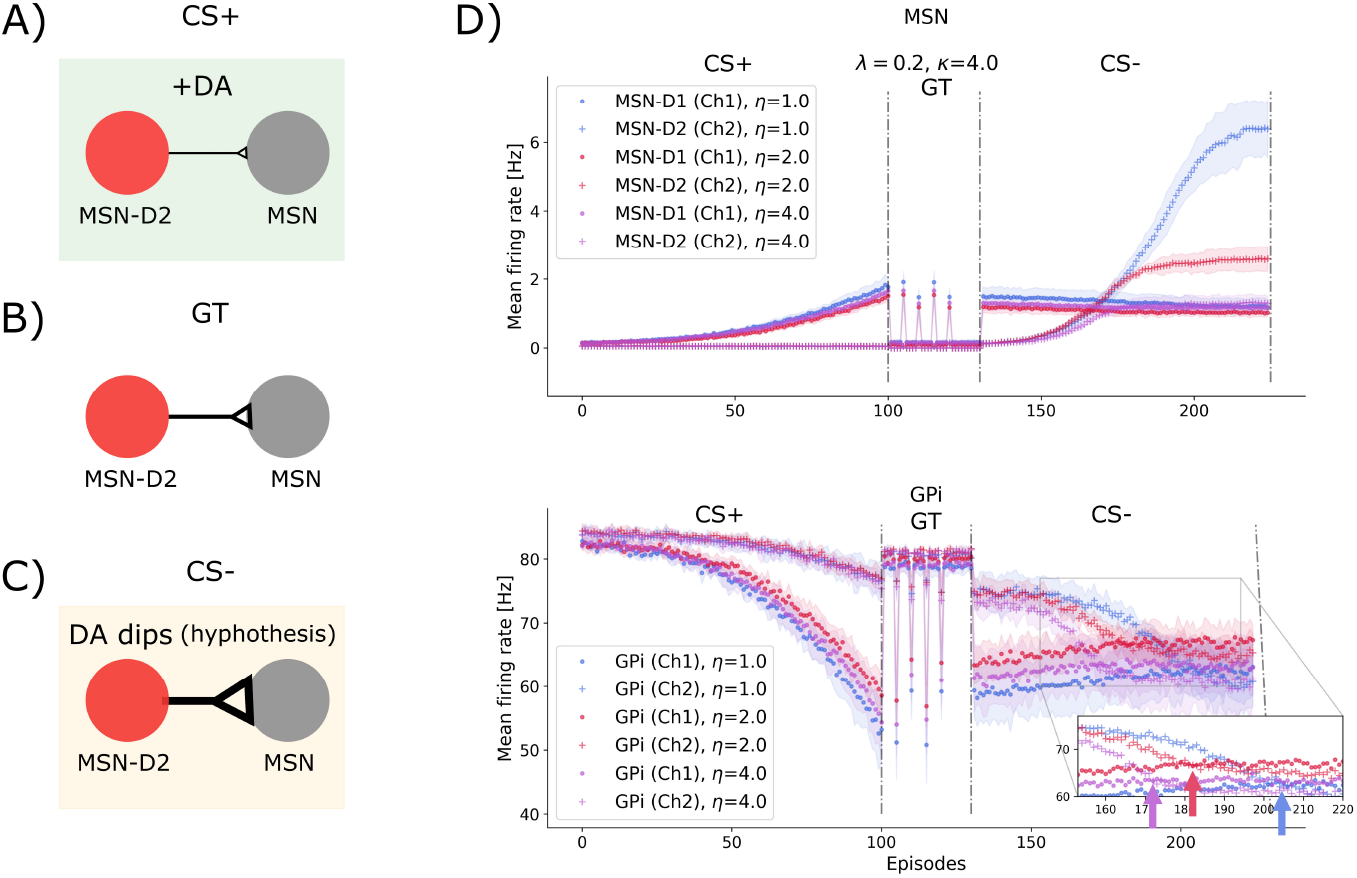
Modulatory effects of DA on axon collaterals. A: DA activation reduces inhibition from MSN-D2 neurons to other MSNs. B: Tonic DA does not modulate collaterals from MSN-D2 neurons. C: DA depletion may enhance inhibition from MSN-D2 neurons (hypothesis). D: Modulation of collaterals is critical for convergence during CS-. A modulation parameter, *η*, enhances collaterals from MSN-D2 neurons during DA depletion, promoting convergence, with stronger modulation resulting in faster convergence (arrows). In the absence of modulation (*η* = 1), convergence is not guaranteed, especially at low *λ* values.

Based on these findings, in the absence of DA changes, that is under tonic DA activity, there is no modulation of the collateral strength. For example, in our simulations, this corresponds to the GT stage, where DA concentration remains constant (Fig.7B).

Following this logic, modulatory effects on MSN-D2 collaterals may also occur during CS-, this time enabled by DA dips. We hypothesize that DA depletion modulates MSN-D2 collaterals, increasing their inhibitory strength toward all MSNs (Fig.7C). This modulation is crucial for achieving convergence in discrimination learning, especially in cases of low pathway overlap.

To test our hypothesis, we conducted systematic simulations with fixed values of *λ* = 0.2 and *κ* = 4 (Fig.7D). In these simulations, we introduced a new parameter, modulation *η*, which is activated only during CS-trials. This parameter enhances the GABAergic transmission of collaterals by a factor of *κ* × *η*.

Without modulation (*η* = 1), convergence slows down, potentially leading to insufficient learning or requiring more trials (see Fig.5 for low *λ* = {0.0, 0.1}), or learning occurring only at the end of the CS-trials (Fig.6C, *λ* = 0.2). This effect can be attributed to the limited number of synaptic connections from MSN-D2s that inhibit the preferred GPi channel 2. However, with *η* = 2, convergence is achieved faster within the 100 CS-trials, and even more rapidly with *η* = 4 (Fig.7D).

It is worth mentioning that we derived this hypothesis from experimental data recorded during DA bursts and speculated about reverse modulation during DA dips. While the predictions of our model provide insight into CS-learning, further evidence is needed to validate this mechanism.

### Overlapping direct-indirect pathways regulate discrimination learning convergence

During CS-trials, the omission of reward and a longer time window for DA dip (0.4s compared to 0.3s for DA burst, see Fig.3A, B) trigger rapid increases in the excitability of MSN-D2s, leading to inhibition of GPi at channel 2.

We observed that different degrees of *λ* directly influence discrimination learning. Specifically, the number of presynaptic connections from MSN-D2 cells providing GABAergic afferents to GPi, as well as the number of presynaptic connections from MSN-D1 cells projecting to GPe, impact the timing of prediction refinement during discrimination learning.

A low value of *λ* implies sparse projections from MSN-D2 to GPi. Our simulation results showed that a certain degree of overlap is required for convergence; otherwise, discrimination learning does not occur (Fig.5A, *λ* = 0.0), or it requires a larger number of episodes (e.g. *λ* = 0.1), resulting in slower changes at GPi channel 2.

In contrast, greater pathway overlap facilitates convergence, resulting in the selection of channel 2 in response to stimuli B. Our simulations reveal diverse convergence trajectories for different *λ* values, as shown in Fig.5A for *λ >*= 0.1. Interestingly, higher *λ* values lead to more rapid convergence during CS-.

To further investigate this effect, we performed systematic simulations with fixed asymmetry *κ* = 4 and varying overlap *λ* = {0.2, 0.3, 0.4} (Fig.6C). Previously, we showed for CS+ that higher degrees of overlap led to stronger disinhibition in GPi channel 1; thus, at the onset of CS-, the gap between the baseline activity of GPi channel 2 and GPi channel 1 is smaller for higher *λ*, promoting faster learning. In addition, increased overlap results in greater inhibition from MSN-D2s to GPi channel 2, accelerating changes in its activity towards that of GPi channel 1. Therefore, the convergence time in discrimination learning is regulated by the overlap parameter *λ* .

### Generalization-discrimination learning is disrupted in a model of Parkinson’s disease

We conducted simulations of a dopamine (DA) imbalanced system, resembling animal models of Parkinson’s disease (PD)^11^, to investigate the impact on learning during a generalization-discrimination task (Fig.8A,B).

**Figure 8.**
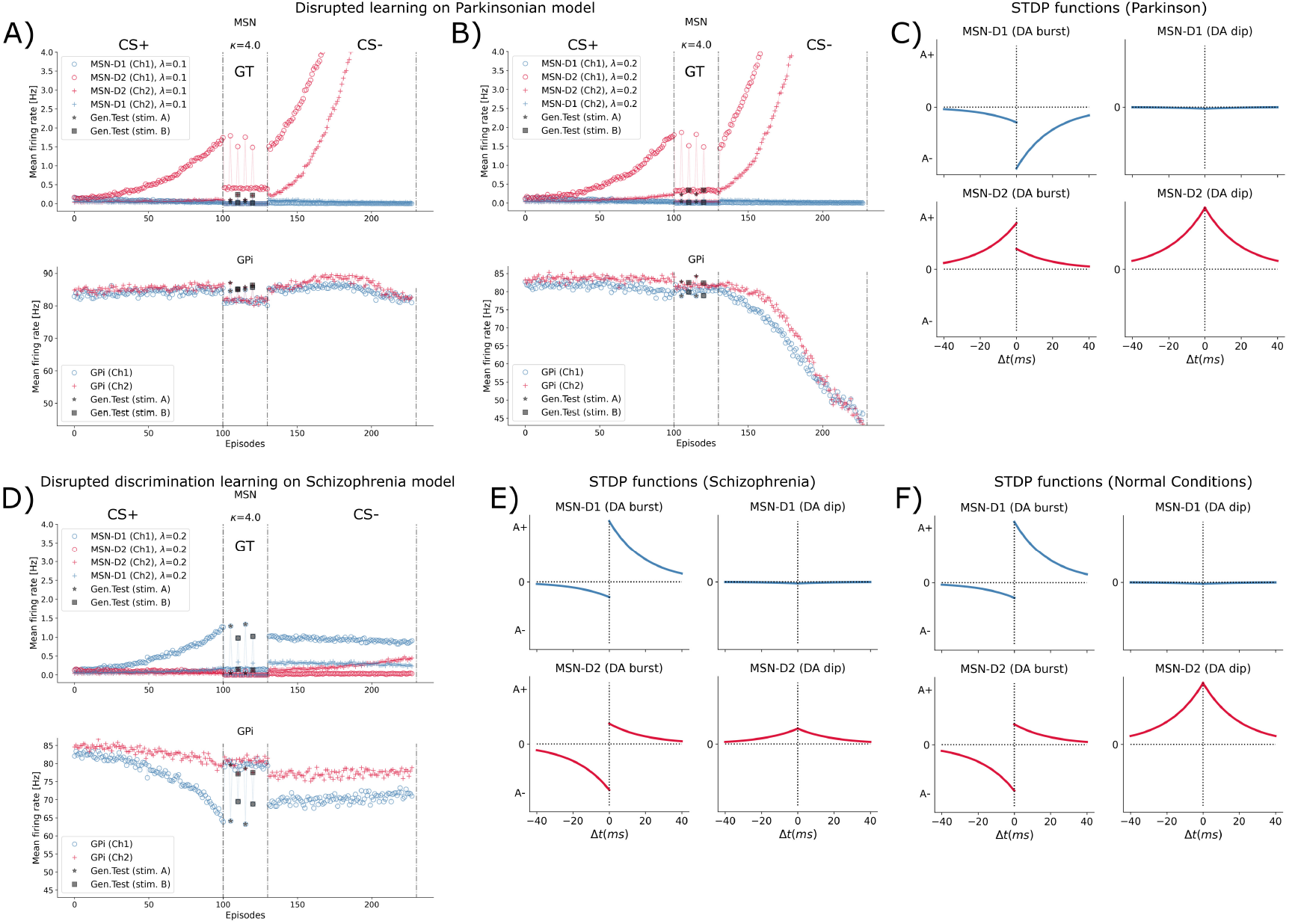
A: Parkinson’s disease model. Learning is simulated with unidirectional STDP changes. CS+ and CS-are both disrupted, and GPi activity shows unclear action selection. B: Effect of pathway overlap (*λ*). Increasing *λ* shows similar learning disruptions: GPi receives strong MSN-D2 inhibition, and action selection remains unclear. C: STDP under hypodopaminergic state. Reduced dopamine (left column) converts MSN-D2 LTD to LTP and MSN-D1 LTP to LTD. D: Schizophrenia disorder model. CS+ learning and the generalization test (GT) are comparable to control, but CS-is impaired: GPi fails to favor channel 2 due to insufficient MSN-D2 activity at channel 2. E: Shortened dopamine dips in CS-. Modeled as a reduced-amplitude STDP function for MSN-D2 during DA dips. F: STDP models under normal conditions.

In animal models of PD, there is a reduction in DA levels, which consequently affects the induction of synaptic plasticity. To examine the effects of DA imbalances during learning, we replicated unidirectional STDP changes observed in previous studies of mouse models of PD^11^, where DA neurons were lesioned or depleted, resulting in a significant reduction of DA levels and an alteration of STDP, leading to a hypodopaminergic state.

In models of PD, although long-term potentiation (LTP) in MSN-D2 neurons remains unaffected, long-term depression (LTD) is lost and instead exhibits changes similar to LTP. Conversely, in MSN-D1 neurons, LTP is absent and instead exhibits changes resembling LTD. Consequently, the bidirectional STDP at glutamatergic synapses from the cortex to MSNs shifts to a unidirectional pattern. In our model and learning task, these changes in DA were implemented by changing the STDP functions for MSN-D1 and MSN-D2 during DA burst (Fig.3D,8F) to unidirectional STDP functions (Fig.8C). For MSN-D1, unidirectional STDP involved reversing the positive amplitude (A+) to negative, while for MSN-D2, the negative amplitude (A-) was reversed to positive.

Our simulations, with fixed values of *κ* = 4 and *λ* ∈ 0.1, 0.2, revealed disrupted conditioning (CS+), manifested as a slow increase in MSN-D1 activity in channel 1 (Fig.8A,B). For *λ* = 0.1, downstream GPi activity showed no clear action selectivity, resulting in exploratory behavior during most CS+ trials. Although synaptic plasticity was present, its dependence on the temporal structure of spiking activity (bidirectionality) was lost, degrading CS+ performance and suggesting perturbed cognitive processes. For *λ* = 0.2, although inhibition of channel 1 in the GPi was slightly stronger due to greater overlap *λ*, action selection remained inefficient.

During subsequent discrimination learning (CS-), STDP increased firing in MSN-D2s at channel 2, yet discrimination did not emerge, indicating perturbation of CS-as well. In the *λ* = 0.1 case, the GPi showed no inhibition, whereas the higher overlap condition *λ* = 0.2 inhibited both channels over episodes, preventing clear discrimination.

Previous reports have suggested that striatal learning is impaired in patients with PD, possibly reflecting reduced cognitive control due to dopamine loss^41^. Consistent with these findings, our simulations show impairments in both CS+ and CS-.

A previous investigation of how PD affects the way people learn from positive versus negative feedback found that unmedicated patients with PD (with low dopamine levels) learned better from negative outcomes (sticks) than from positive outcomes (carrots)^42^, while the same patients on dopamine medication showed the opposite pattern, learning better from positive feedback and relatively worse from negative feedback. In other words, off-medication PD patients excelled at avoiding choices that led to bad outcomes, but struggled to learn from rewards, while dopamine replacement therapy reversed this bias. While our model reproduced impaired reward/positive learning (CS+), consistent with off-medication PD showing poorer carrot learning, our simulations do not show the enhanced negative/avoidance learning (CS-) seen off-medication; instead, CS-is also impaired. This points to a global selection/learning deficit rather than a valence-specific bias.

### Discrimination learning is disrupted in a model of schizophrenia disorder

Previous experiments with methamphetamine (MPA) treatment in mice showed that dysregulated DA signaling shortened the DA dips in MSN-D2 neurons observed during discrimination learning^13^. This impairment affected discrimination learning and is believed to contribute to neuropsychiatric symptoms associated with D2 receptors, such as those seen in schizophrenia.

To investigate this, we modeled schizophrenia by modifying the STDP function for MSN-D2 during DA dips in our model. Specifically, we reduced the amplitude (A+) of the STDP function to 1*/*4 of its original value (Fig.8E) and tested its impact on generalization-discrimination learning.

In our simulations with fixed values of *κ* = 4 and *λ* = 0.2, CS+ is not affected, since the simulations involve STDP functions for the DA burst. Similarly, there is no observed effect on GT. However, during CS-phase, where STDP for DA dips is used, exposure to MPA and the resulting increase in DA concentration are simulated by applying the previously mentioned STDP update.

As a result, while CS+ and GT outcomes align with previous simulations under normal conditions (Fig. 4), CS-appears impaired (Fig.8D). The reduced DA dips hindered learning during CS-, leading to the persistent selection of GPi channel 1 (the CS+ preferred action) while GPi channel 2 did not receive strong inhibition from MSN-D2s. Consequently, it maintained steady activity over the 100 CS-episodes, preventing convergence.

### Large-scale simulations

Smaller-scale models are useful for implementation and analysis, but their reduced network sizes have inherent limitations, as highlighted in previous studies^43^. Large-scale brain simulations are vital for uncovering complex neural interactions, modeling brain disorders, and safely testing hypotheses.

We simulated different biological scales in our model by adjusting the reference patch area (Fig.9A), which increased the number of neurons and connections. Table 1 outlines the scaling of the reference patch size and its equivalencies with real-world brain sizes, covering both hemispheres and single hemispheres across species. Assuming the basal ganglia (BG) account for approximately 3% of the total neurons in the brain, our simulations represent the entire BG of rodents^44^ and the BG of a single hemisphere for marmosets and galago monkeys^45^.

**Figure 9.**
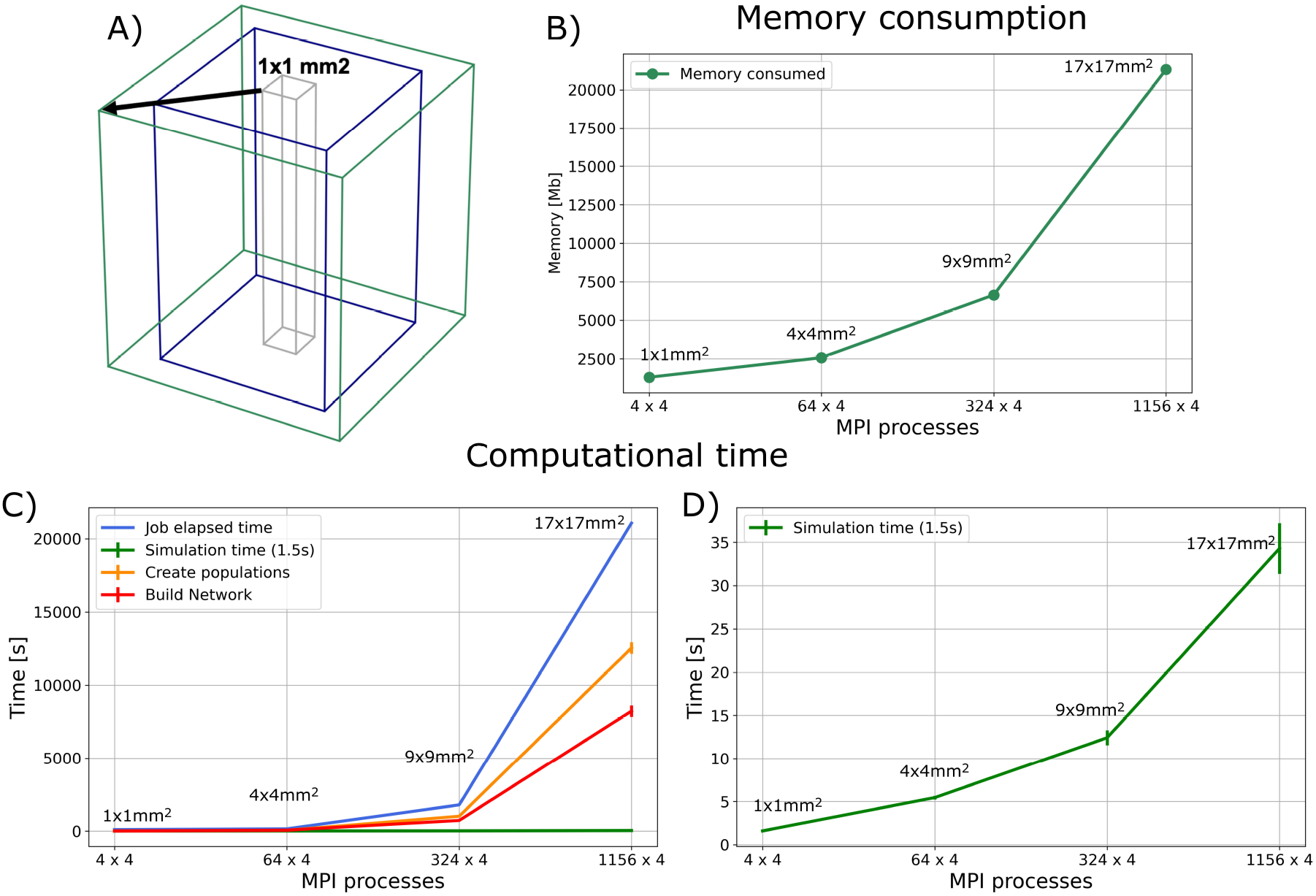
Simulation performance on Fugaku: A) Schematic representation of scaling by increasing the size of the reference patch for the nuclei. B) Memory consumption increases with the construction of larger networks. Computational resources are indicated by the number of MPI processes, with 1,156 nodes used for the largest network (each node comprising 4 processes and 12 threads). C) Network construction time increased proportionally with the size of the basal ganglia (BG) model and the computational resources. D) Simulating 1.5 seconds of biological time showed efficient performance, with only minor increases in simulation time for larger models.

**Table 1.**
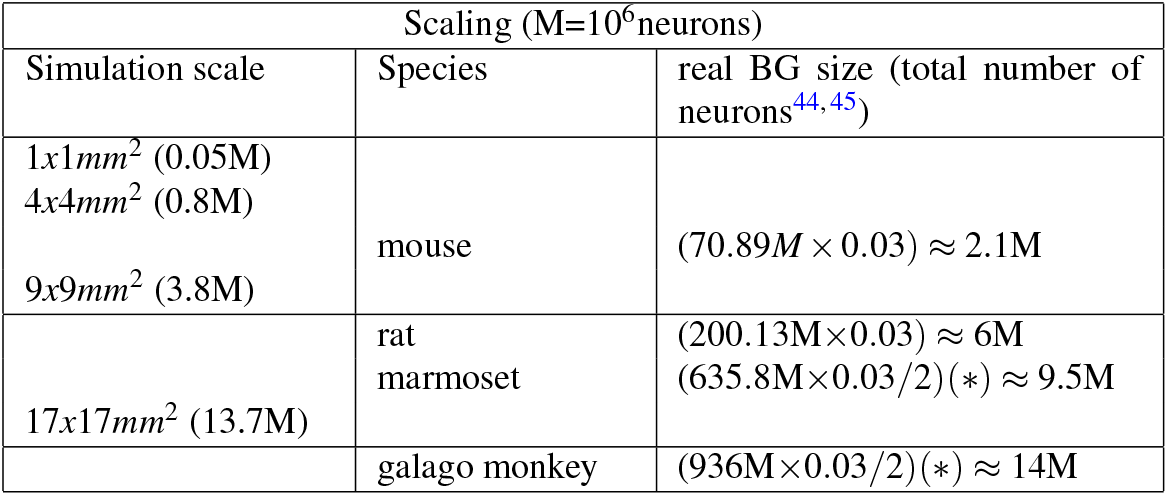
The BG model was scaled across a range of sizes, from 1×1*mm*^2^ to 17×17*mm*^2^ of the reference patch, representing realistic BG sizes in rodents and single-hemisphere sizes in non-human primates (*).

Simulations were carried out on Fugaku, Japan’s next-generation flagship supercomputer^46^, using a weak-scaling scheme. This approach proportionally increases both the network size and the number of computational nodes (MPI processes). A hybrid parallelization method was applied, combining multiple MPI processes with 12 OpenMP (https://www.openmp.org/) threads per MPI. Scaling performance was assessed based on network construction time, simulation runtime, and memory usage (Fig.9).

At the largest simulated scale, total memory usage (network construction and simulation) reached approximately 20GB per computational node, with most of it consumed during network construction (Fig.9B). Network construction required about 5.5 hours (20,000 seconds) at this scale (Fig.9C), while simulating 1.5 biological seconds took only 35 seconds (Fig.9D), demonstrating efficient scaling performance for the simulation.

## Discussion

We have presented a topologically organized model of the macaque basal ganglia, with striatal symmetries, channel formation, overlapping direct and indirect pathways, and dopamine-enabled STDP. The model performed resting state and action selection simulations.

In the action selection simulations, neighboring channels received overlapping signals from cortical afferents. The results showed relatively high efficiency but also high distortion due to interference from neighboring channels, while non-neighboring channels exhibited high efficiency in channel selection and less distortion. This can be assumed and used to study the competition between two similar inputs (e.g., similar sounds at close frequencies), which could be allocated to neighboring channels, and dissimilar inputs (e.g., sound and vision), which could be allocated to non-adjacent channels.

We investigated, through simulation, the transient increases and decreases of dopamine in generalization-discrimination learning, following the results from Iino, Yagishita, and colleagues^12,13^. During simulations, we analyzed the effect of two parameters: the degree of strength-dependent asymmetry from MSN-D2s to other MSNs (denoted as *κ*) and the degree of overlap between direct and indirect pathways (*λ*). Our results revealed that overlapping direct-indirect pathways decrease action-selection efficiency in conditioning learning, and that efficiency decreases further with higher *λ* . This could indicate that a certain overlap between these pathways may disrupt the balance required for efficient action selection, possibly leading to signal degradation through dispersion in the pallidum.

We also observed that lateral inhibition from MSN-D2s (*κ*) in discrimination learning caused a gradual and slight decrease in MSN-D1 activity. These collaterals contribute to downstream disinhibition of the GPi, which is also noticeable when the pathways are discriminated (low *λ*). This is thought to support discrimination learning but does not guarantee learning convergence, especially for low overlap.

Moreover, we further modulated these collaterals enabled by dopamine dips with a parameter *η* × *κ*. This modulation was necessary for convergence, adjusting the firing rate of MSN-D1s. This modulation, *η*, corresponds to the hypothesis that dopamine depletion modulates MSN-D2 collaterals, increasing their inhibitory strength toward all MSNs, and is crucial for convergence in discrimination learning. This hypothesis is derived from experimental observations of the depression of MSN-D2 collaterals during dopamine activation. First, these findings highlight the importance of intra-striatal interactions in modulating action selection, with MSN-D2s “fine-tuning” MSN-D1 activity and ensuring that competing actions are constrained. Secondly, while the *κ* mechanism facilitates discrimination, it alone may not be sufficient for robust action selection. Our results suggest that dopamine depletion enhances MSN-D2 collateral inhibitory strength. The balance between dopamine-dip-mediated modulation and lateral inhibition seems to be a key determinant of efficient discrimination learning. Dysregulation in either process could lead to impaired learning.

Furthermore, we found that direct-indirect pathways may also regulate discrimination learning convergence. We observed that different degrees of *λ* directly influence discrimination learning. Insufficient overlap impairs learning, and a certain degree of overlap is required for convergence, with diverse convergence schedules for different *λ* . These results challenge the notion of perfectly segregated pathways and highlight the potential functional role of overlapping pathways. Some overlap may be critical for functional adaptability during learning, and we speculate that *λ* might be related to the learning capabilities of the subject or species. High levels of overlap were observed in non-human primates, while rodents exhibited a lower degree of overlap.

Thus, overlap could correlate with the complexity or efficiency of learning mechanisms and implies that *λ* might serve as a neuroanatomical marker or connectivity feature of learning potential across species. The variability in overlap might reflect evolutionary adaptations, where higher *λ* facilitates more sophisticated or flexible learning in species with complex behavioral repertoires, such as primates. Speculating on the generalization of our basal ganglia model predictions to the human brain, it is possible that the human basal ganglia exhibit a high degree of overlap between direct and indirect pathways. Few studies have investigated the association between the basal ganglia and intelligence in humans. However, recent results based on MRI imaging and sub-cortical volume metrics suggest the importance of the basal ganglia in intelligence, cognitive flexibility, and working memory^47^. These findings highlight the importance of interactions between direct and indirect pathways in regulating learning efficiency and convergence, suggesting a broader role for these pathways beyond action selection.

In addition, we have implemented Parkinson’s disease and schizophrenia disorder and studied by simulation their impact on generalization-discrimination learning. Simulations with the modeled unidirectional STDP functions for Parkinson’s disease revealed disrupted conditioning with slow increases in MSN-D1 activity but no clear action selectivity, suggesting perturbed learning. Discrimination learning did not occur, although there were increases in MSN-D2 activity. Simulations reveal that dopaminergic dysregulation in Parkinson’s disease leads to impaired synaptic plasticity, likely affecting the basal ganglia’s ability to adapt to new learning experiences.

Similarly, simulations with the reduced amplitude of the STDP function for MSN-D2 during dopamine dips to mirror dopaminergic dysregulation observed in schizophrenia disorder showed that while conditioning remained unaffected, discrimination learning was impaired. The inability to correctly perform discrimination learning despite normal conditioning suggests that action selection might still occur, but the ability to prioritize or inhibit less appropriate actions is impaired, which aligns with the cognitive deficits observed in schizophrenia. Conditioning relies more on the direct pathway (MSN-D1), which is less affected by changes in dopamine dips. However, discrimination learning likely involves more complex integration of both pathways, and dysfunction in MSN-D2 during dopamine dips impairs this process.

Finally, we have performed large-scale simulations on the Fugaku supercomputer for resting state, showing promising performance for future human-size basal ganglia simulations. Moreover, considering integration with the Brain/Minds 2.0 project^48^, there is a potential need for neural simulations at realistic scales for the marmoset brain. A full marmoset brain consists of approximately 635 million neurons^45^. Thus, 3% of a single hemisphere would encompass around 9.5 million neurons for the basal ganglia. We successfully generated a comparable model of the basal ganglia for the marmoset using relatively few computational resources (1,156 nodes from a total of 158,976 available nodes). This demonstrates the feasibility of non-human primate-scale and human-scale brain simulations.

In conclusion, the interplay between collateral dynamics, lateral inhibition, overlapping pathways, and dopamine signaling underscores the intricate mechanisms underlying basal ganglia-mediated learning. It also suggests that fine-tuning inhibitory and excitatory interactions is vital for adaptive behavior, with potential implications for both normal brain function and neurological disorders.

## Methods

### Model parameters and topological organization

The model incorporates a set of parameters described in Girard et al.^7^, with adjustments made to accommodate the new 2D topographical organization based on a reference patch corresponding to the model scale.

#### Number of cells and scale

The number of cells, *N*, is based on previous efforts^6,7^, where the number of MSNs was derived from cell counting in 0.01*mm*^2^ patches of frozen 30*μm* sections from rhesus monkey^49^. For simulation purposes, the model scales increases the number of 0.01*mm*^2^ patches and the corresponding *N* proportionally across the nuclei.

A reference patch scale of *S* × *Smm*^2^, where *S* = 1.0 ∼ 1.2*mm*, provides a reasonable number of neurons for the sake of experimentation by simulation. The selected size of the BG model represents approximately a scale of 1*/*10000 × 4 × 3 (1:834) of the real number of neurons found in the macaque BG^6^ (see Table.2) for details of *N*.

The scaled tissue is modeled as a 2-dimensional layer, with a single layer per nucleus. The spatial organization of neurons is approximated by a random-uniform allocation within the reference area.

This scale allows, for example, the inclusion of approximately 100 cells in the nucleus with the fewest cells, the STN. Similarly, in the output nuclei GPi/SNr, where cell density is much lower compared to the input layer (MSN), around 170 cells were simulated (Fig.1D). These numbers are reasonable, as the columnar organization is crucial, and each column requires a few neurons in the STN and GPi/SNr to enable meaningful experimentation in the simulations.

#### MSNs organization

The model assumes a generic striatum with no distinction between matrix and striosome^50^. MSN neurons were organized in two subtypes^8,51^, based on their expression of dopamine receptors, 50% are MSN-D1 (dopamine D1 receptors) and the other half MSN-D2 (dopamine D2 receptors).

To ensure equally distributed MSN types, a single random-uniform position is shared by one MSN-D1 and one MSN-D2. Thus, allocation of neurons near the boundaries is avoided by neglecting 0.1*mm* from the left and right (up and down) of the 2D layer. The resulting spacing of neurons for 1:834 scale is assumed biologically plausible for the sake of simulations.

Fig.10A illustrates the various inputs converging onto the striatum, including PTN and CSN from the cortex, and CMPf from the thalamus. The number of converging connections is shown for different quantities of MSNs, ranging from a single neuron to clusters of MSNs arranged in square patches with side lengths of 0.09, 0.15, and 0.3*mm*. Asymmetries in the striatum were modeled in terms of the proportions of connections between MSNs and their connection weights, as detailed in Section 2 of the Results above.

**Figure 10.**
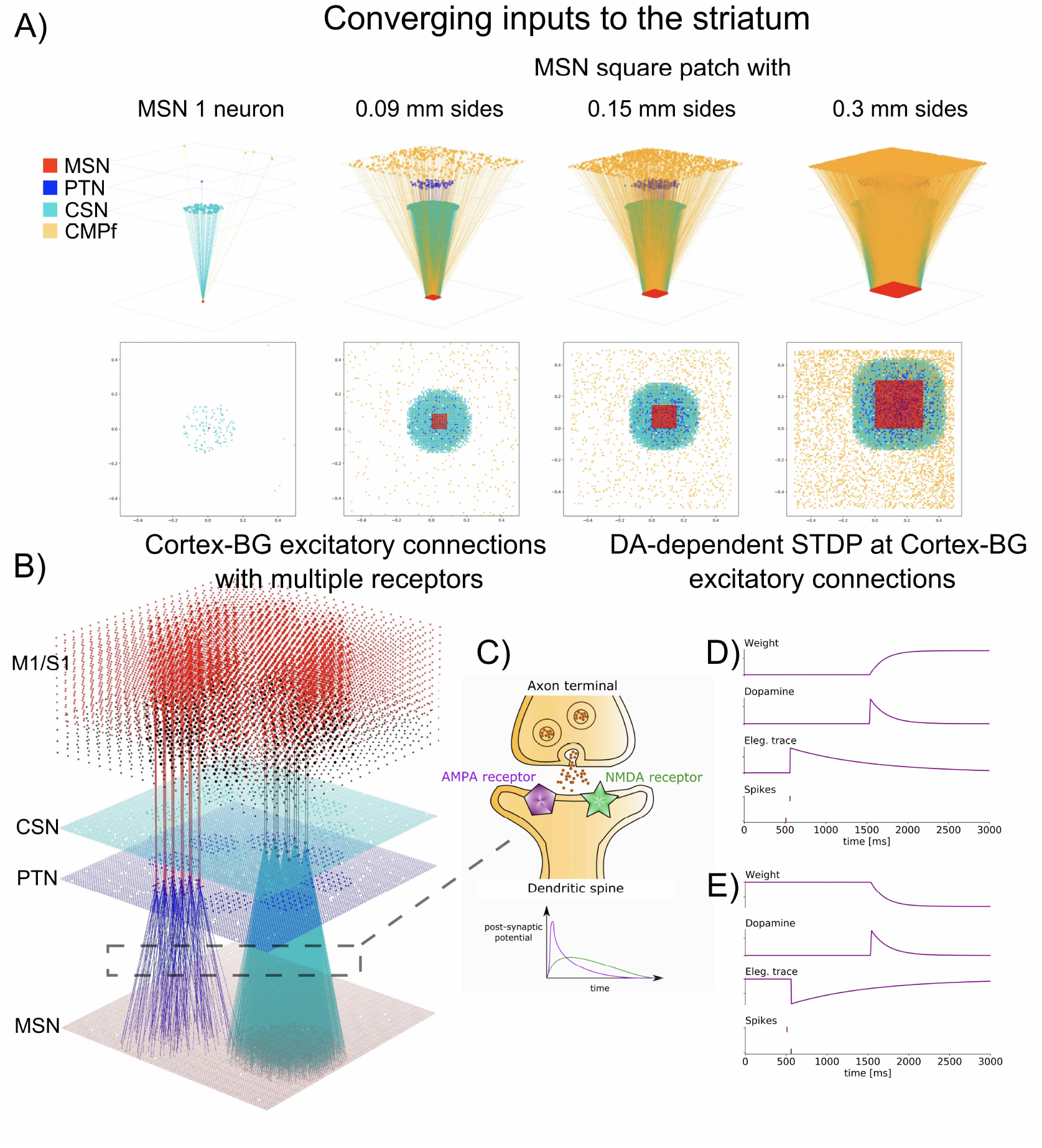
A: Topological and topographical organization of basal ganglia (BG) input. PTNs and CSNs form focused converging connections to the striatum, while CMPfs establish diffuse connections, illustrated for different MSNs numbers. B: Projections from motor and sensory regions to the striatum preserve topographical mapping, with adjacent cortical regions projecting to adjacent areas in the striatum. C: All excitatory synapses in the model are mediated by AMPA and NMDA receptors, while inhibitory synapses are mediated by a single GABA receptor. D: The STDP model for excitatory synapses from the cortex to the striatum incorporates dynamics for the eligibility trace, dopamine concentration, and synaptic weight changes, enabling long-term potentiation (LTP, panel D) and long-term depression (LTD, panel E).

#### Neuron parameters

Neuron parameters, including PSP (post-synaptic potential) amplitudes (*V*_*n*_) and their half-times 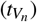, dendritic extent (*l*), and diameter (*d*), were adopted from previous models^6,7^ as fixed values.

The resting potential of all neurons was set to 0*mV*, a neutral shift that does not affect model dynamics. Membrane time constants, firing thresholds, refractory periods, and other parameters were specified according to the optimized values and assumptions of these prior efforts.

Each nucleus received a fixed tonic input, reflecting observed basal ganglia (BG) activity even in the absence of external inputs. This was implemented as constant external currents (*Ie*) using optimized values from earlier work^7^.

Thus, the topologically organized BG model maintains the original parameterization derived from evolutionary optimization based on biological and physiological objectives. For more details, refer to Tables 1 and 2 in^7^ and^6^.

**Table 2.**
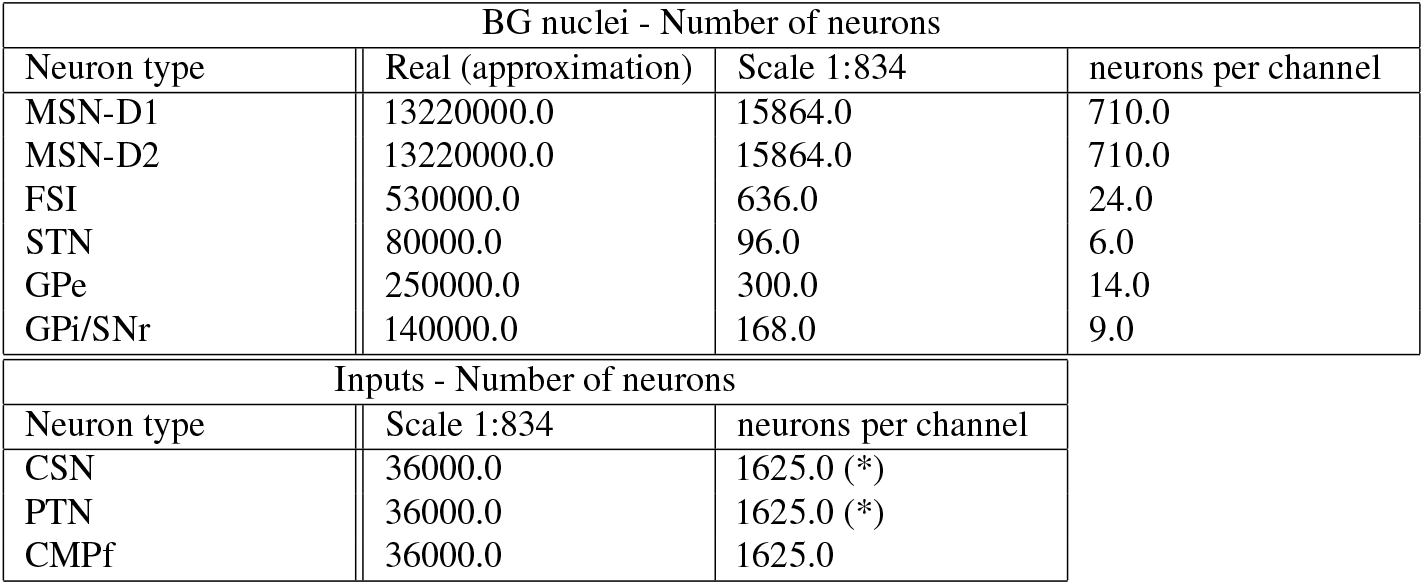
Number of neurons: real BG size and 1:834 model-scale numbers. Channels allocates a cluster of neurons within a circular area of 240m*μ* diameter. (*) For channels simulation experiments, a set of 200 ∼ 300 neurons from CSN and PTN each were randomly selected and stimulated.

#### Inputs and connectivity

To model the cortical contribution to the formation of discrete functional units (channels) processing segregated information streams in the basal ganglia (BG)^52^, cortical afferents to the striatum were designed as projections with sufficient spatial spread. This configuration ensured that nearby MSNs received shared cortical inputs^53^, while MSNs located near or between channels likely integrated inputs from distinct cortical activity patterns^54^ (overlapping inputs). At the same time, topographic mapping was preserved, with neighboring cortical regions projecting to adjacent striatal regions^52,55^ (Fig.1E, 10A,B).

The spatial spread was implemented using a circular mask with a 0.15*mm* radius. Each MSN was placed at the center of the mask, and incoming connections were randomly selected from source nuclei (PTN and CSN) within the mask’s bounds.

For thalamic input (CMPf), a diffuse connection pattern was modeled using a larger circular mask with a 2.0*mm* radius. This approach reflected experimental findings that thalamic axons exhibit a diffuse organization across the striatum^6^ (Fig.10A). Additional details about connectivity are provided in Table 3.

**Table 3.** Connectivity Scheme based on^6^.

| BG nuclei connectivity (source→target) |  |
| --- | --- |
| Focused-type<br>(0.15mm radius<br>circular mask) | Diffuse-type (2.0mm<br>radius circular mask) |
| CSN→MSN<br>PTN→MSN<br>MSN→MSN | CMPf→MSN<br>FSI→MSN<br>STN→MSN<br>GPe→MSN |
| CSN→FSI<br>PTN→FSI | CMPf→FSI<br>FSI→FSI<br>GPe→FSI<br>STN→FSI |
| PTN→STN<br>GPe→STN | CMPf→STN |
| MSN→GPe | CMPf→GPe<br>STN→GPe<br>GPe→GPe |
| MSN→GPi<br>GPe→GPi | CMPf→GPi<br>STN→GPi |

#### BG Nuclei connectivity

Similarly, the connectivity between BG nuclei incorporates two distinct spatial spreads: a focused connection, defined by a circular mask with a radius of 0.15*mm*, and a diffuse connection, characterized by a circular mask with a radius of 2*mm*. While the focused spread allows for some degree of synaptic input overlap, the diffuse spread results in highly overlapping connections across the entire surface of the reference tissue patch.

These connectivity schemes are adapted from the approach in Lienard and colleagues^6^, with the addition of overlapping input features introduced by the topographic organization of the current model. Table 3 outlines the connectivity types for each population pair.

The number of incoming synapses per neuron, defined by the in-degree parameter, was previously optimized^7^ to align with plausible values of synaptic contacts or bouton counts. The model implements all the excitatory glutamatergic synapses mediated by AMPA and NMDA receptors (Fig.10C), inhibitory GABAergic signals through a single GABA receptor, and dopaminergic signals via D1 and D2 receptors, specific to MSN-D1 and MSN-D2 neurons, respectively.

#### Neuron model

We modeled all neurons as point neurons of the Leaky Integrate-and-Fire (LIF) type with multi-synapses. The neuronal dynamics for each BG neuron type were designed as an integration process combined with a mechanism for generating action potentials when the membrane potential crosses a threshold *ϑ* . The membrane potential *u*_*i*_(*t*) reflects the cumulative effect of all synaptic inputs. When *u*_*i*_(*t*) reaches *ϑ* from below, the neuron *i* fires a spike at time 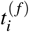. In the LIF model, action potentials are simplified as discrete events^56^.

The LIF dynamics consist of two components:

i. Membrane potential evolution: 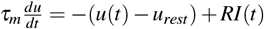, where *u*(*t*) is the membrane potential, *u*_*rest*_ is the resting potential, *I*(*t*) is the input current, *R* is the leak resistance, and *τ*_*m*_ is the membrane time constant.
ii. Spike generation and reset: When *u*(*t*) *> ϑ*, a spike is emitted, and *u*(*t*) is reset to a defined value.

For implementation, we used the NEST simulator’s multi-synapse LIF model, `iaf_psc_alpha_multisynapse`, to represent the different neuron types in the BG model.

#### DA-enabled STDP

In our model, dopamine is considered to influence the presynaptic terminals of cortical inputs to the striatum, reinforcing D1 receptor activity in MSN-D1 neurons during DA bursts and D2 receptor activity in MSN-D2 neurons during DA dips, as reported by Iino et al.^13^. These dynamics result in long-term changes in synaptic strength (Fig.10D,E).

We incorporated spike-timing-dependent plasticity (STDP) at NMDA and AMPA excitatory synapses from the cortex to the basal ganglia (BG) using the model from^57,58^ (Eq.1, 2). The eligibility trace and dopamine concentration dynamics govern long-term potentiation (LTP, Fig.10D) and long-term depression (LTD, Fig.10E) based on the inter-spike interval. The mathematical model for synaptic weight changes, *w*, is expressed in Eq.1, where:

*c* represents the eligibility trace, *n* is the neuromodulator concentration, *s*_*pre/post*_ denotes pre-/post-synaptic spike times, *s*_*n*_ represents neuromodulatory spike times, *b* = 0 is the baseline dopaminergic concentration, *C*_1_ = 1 and *C*_2_ = 1 are constants, *τ*_*c*_ = 700*ms* (eligibility trace time constant) was tuned to align with the critical time window of ∼ 2*s*^13^ (Fig.3A, B, E), *τ*_*n*_ = 100*ms* (dopaminergic trace time constant) was set based on previous studies^59,60^.

The dopamine-modulated STDP update rule (Eq.2) uses Δ_*t*_, the temporal difference between post-synaptic and pre-synaptic spikes, a single STDP time constant (*τ*_*stdp*_ = 20*ms*) which governs facilitation and depression, and *A*_+_ and *A*_−_ representing the amplitudes of weight changes. These amplitudes, which depend on the sequence of post- and pre-synaptic spikes, were scaled proportionally to the values in^61^, derived from in vitro data^11^ for MSN-D1 and MSN-D2 under high and low dopamine levels (Fig.3D).

For implementation, we used the `stdp_dopamine_synapse` model^57,58^ from the NEST simulator (version 2.20)^39^. This model integrates volume transmitter devices to diffusively deliver spikes from dopaminergic neurons to cortical-striatal synapses. Dopaminergic neurons were modeled as a Poisson process emitting spikes at 20Hz to feed the volume transmitters.

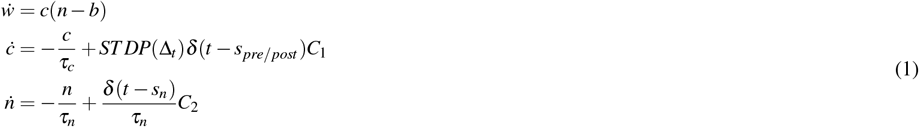

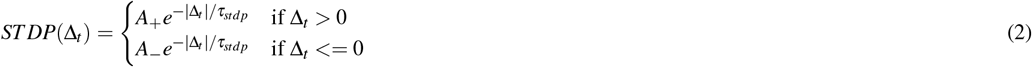

## Code availability

The code will be made publicly available on GitHub once the paper is accepted.

## Acknowledgements

This research was supported by the Brain/MINDS 2.0 Program, the Digital Brain, from the Japan Agency for Medical Research and Development AMED. The computational nodes of the Fugaku supercomputer were provided by the Next Fugaku brain simulation project.

## Author contributions statement

Conceptualization, C.G. and K.D.; model development, C.G. and J.L.; learning implementation, C.G.; mapping to generalization–discrimination learning, advised by H.U.; basal ganglia (BG) modeling, validation and experimental / simulation design, advised by B.G. and K.D.; all authors wrote and reviewed the manuscript.

